# Histidine-rich protein 2: a new pathogenic factor of *Plasmodium falciparum* malaria

**DOI:** 10.1101/2021.11.19.469193

**Authors:** Takashi Iwasaki, Mayu Shimoda, Haru Kanayama, Tsuyoshi Kawano

## Abstract

*Plasmodium falciparum* causes serious malaria symptoms; when this protozoan parasite infects human erythrocytes, it produces and secretes large amounts of histidine-rich protein 2 (PfHRP2) into human blood. Thus, PfHRP2 is a well-known diagnostic marker for malaria infection. Here, however, we also identified PfHRP2 as a pathogenic factor produced by *P. falciparum*. PfHRP2 showed cell penetration and cytotoxicity against various human cells. In particular, PfHRP2 showed significant cytotoxicity over 5 days at the same concentration as in *P. falciparum*-infected patients’ blood (90–100 nM). This result is consistent with the mortality rate of *P. falciparum* malaria, which increases rapidly in untreated cases for 3–7 days. In addition, the cell penetration and cytotoxicity of PfHRP2 increased 2.5- and 2.6-fold, respectively, in the absence of serum, which suggests that low serum protein concentrations (occurring during malnutrition, for example) increase the risk of adverse effects from PfHRP2 (consistent with malnutrition increasing the lethality of malaria infection). We also showed that PfHRP2 bound to Ca^2+^ ions, localized to intracellular lysosomes, increased lysosomal Ca^2+^ levels, and inhibited the basal level of autophagy by inhibiting autolysosome formation. Furthermore, the Ca^2+^-dependent cytotoxicity of PfHRP2 was suppressed by the metal ion chelator ethylenediaminetetraacetic acid (EDTA). In summary, our findings suggest that PfHRP2 acts as a pathogenic factor in *P. falciparum*-infected patients and is associated with the exacerbation of malaria. Furthermore, EDTA is a promising candidate as a therapeutic agent for the suppression of PfHRP2 pathogenicity. Overall, this study provides new insights into *P. falciparum* malaria pathogenesis and treatment.

## Introduction

The cell membrane acts as a physical barrier that limits the traffic of hydrophilic biomolecules inside and outside of the cell. In particular, typical hydrophilic biomolecules, such as peptides and proteins, are tightly controlled in terms of intracellular and extracellular localization. However, certain peptides and proteins are selectively taken up by the cell via endocytosis. In our previous studies, we reported a new cell-penetrating synthetic polyhistidine peptide consisting of only 16 histidine residues (1) that is efficiently internalized to various human cells. We also reported that nonmembrane permeable proteins, such as green fluorescent protein (GFP) and the hydrolytic enzyme a-galactosidase A (GLA), fused to the polyhistidine peptide by covalent or noncovalent bonds are similarly internalized to human cells (2). These reports indicate that the artificial histidine-rich sequences in the proteins significantly increase their membrane permeability. This suggests the possibility that natural histidine-rich proteins may also permeate the cell membrane to become internalized into human cells.

By searching for natural histidine-rich proteins via Protein BLAST (https://blast.ncbi.nlm.nih.gov/Blast.cgi?PAGE=Proteins), we identified histidine-rich protein 2 (PfHRP2) from the protozoan parasite *Plasmodium falciparum*. The mature PfHRP2 protein produced by *P. falciparum* consists of 278 amino acid residues with 99 histidine amino acids (35.6% of the total amino acids) (Fig. 1*A*) (3, 4). *Plasmodium falciparum* malaria is an infectious disease in which the pathogenic *Plasmodium* protozoa are introduced into the human bloodstream through the bite of a female *Anopheles* mosquito (5). *Plasmodium falciparum* infects human erythrocytes and induces potentially fatal exacerbation of symptoms including encephalopathy, renal failure, liver disorders, pulmonary edema, severe anemia, metabolic acidosis, hypoglycemia, and black water fever (6). Worldwide, five types of *Plasmodium* species (*Plasmodium vivax, Plasmodium malariae, Plasmodium ovale, Plasmodium knowlesi*, and *P. falciparum*) infect 229 million people a year resulting in 409,000 deaths; however, *P. falciparum* is the deadliest parasite, accounting for nearly 99% of cases in Africa and 94% of all malaria cases in 2019 (7, 8). PfHRP2 is produced by *P. falciparum* and secreted into human blood in large amounts when the parasite infects human erythrocytes (9, 10). Therefore, PfHRP2 is a well-known diagnostic marker for the detection of *P. falciparum* malaria infection (11). The frequent contact between PfHRP2 and human cells suggests that the former may be internalized into the latter in *P. falciparum*-infected patients.

**Fig. 1.**
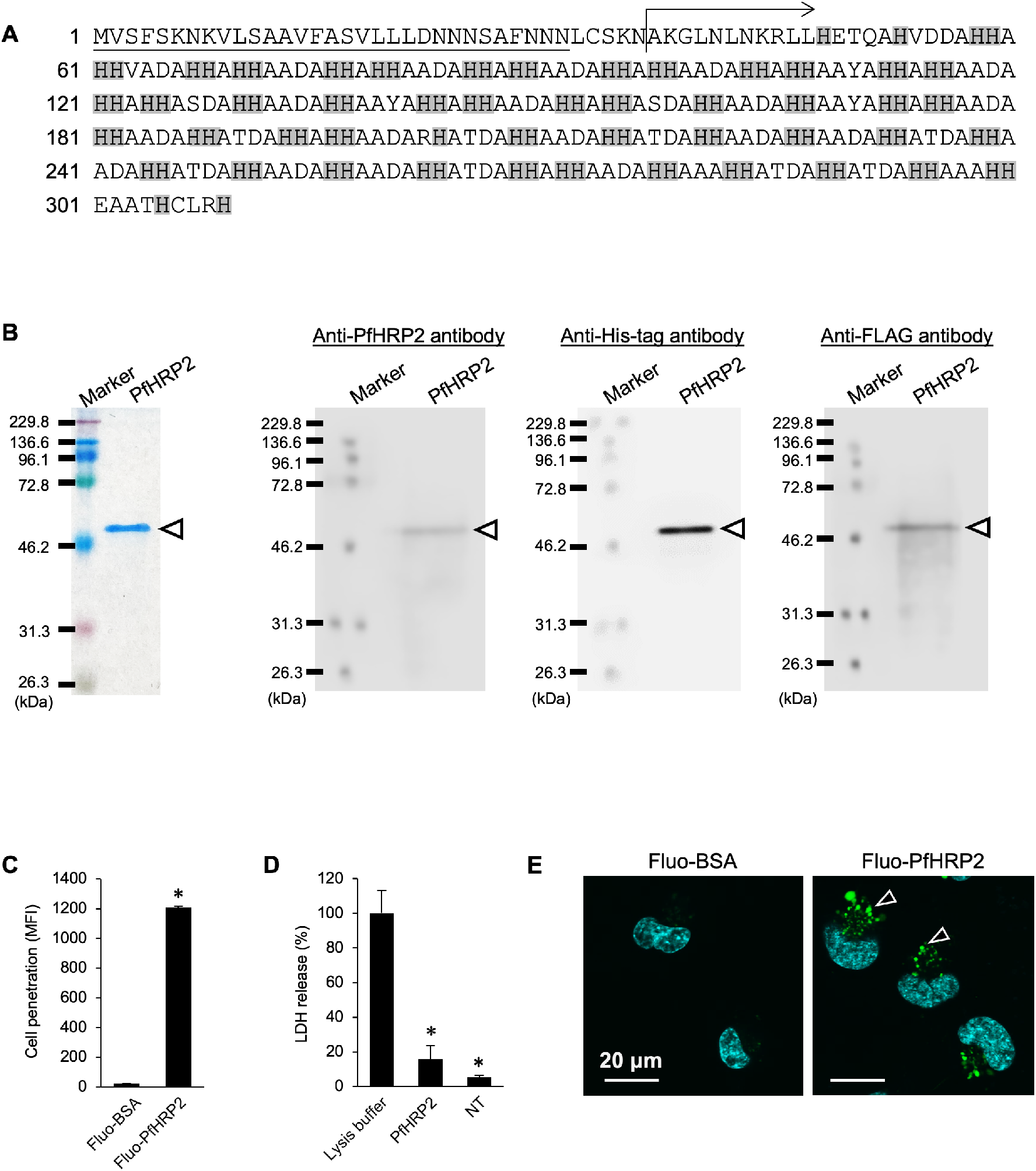
Preparation of recombinant PfHRP2 and its cellular uptake against human fibrosarcoma HT1080 cells. (*A*) The amino acid sequence of *Plasmodium falciparum* histidine-rich protein II (PfHRP2). Underlining and gray boxes indicate the signal sequence and histidine residues, respectively. Arrow indicates initial sequences of recombinant PfHRP2 used in the present study. (*B*) SDS-PAGE and western blotting images of recombinant PfHRP2. The recombinant PfHRP2 with C-terminal FLAG tag (DYKDDDDK) was expressed in *E. coli* and purified using an anti-FLAG affinity column. Purified PfHRP2 was confirmed by SDS-PAGE and western blotting using anti-PfHRP2, anti-His-tag, and anti-FLAG antibodies. (*C*) Cellular uptake of PfHRP2 in human fibrosarcoma HT1080 cells. The HT1080 cells were treated with fluorescein-labeled PfHRP2 and bovine serum albumin (Fluo-PfHRP2 and Fluo-BSA) at 1 μM and 37°C for 3 h. Cell penetration of proteins was evaluated by the mean fluorescent intensity (MFI) in the flow cytometric analysis. Means and SD are shown (\**p* < 0.05). (*D*) Lactate dehydrogenase (LDH) leakage in HT1080 cells. The HT1080 cells were treated with PfHRP2 (1 μM) cells at 37°C for 3 h and then LDH release was determined with a Cytotoxicity LDH Assay Kit-WST according to manufacturer’s protocol. NT indicates nontreated cells used as a negative control. Means and SD are shown (\**p* < 0.05). (*E*) Cellular uptake image of PfHRP2 in HT1080 cells. HT1080 cells were treated with Fluo-PfHRP2 (1 μM) or Fluo-BSA (1 μM) at 37°C for 3 h. Cyan fluorescence and arrowheads indicate nuclei stained with Hoechst 33258 and Fluo-PfHRP2 internalized into HT1080 cells, respectively. Scale bars: 20 μm.

In the present study, we examined the possibility of cell penetration by PfHRP2 in human cells; as expected, recombinant PfHRP2 showed significant cell penetration abilities in various human cells. This finding represents a novel phenomenon that had yet to be reported until now. Furthermore, our experiments showed that PfHRP2 is cytotoxic: it inhibited basal-level autophagy after penetration into human cells. Under the same concentration as that in the *P. falciparum*-infected blood of patients (90–100 nM) (12), PfHRP2 showed significant cytotoxicity, which strongly suggests that PfHRP2 cytotoxicity occurs in nature. Further investigations showed that Ca^2+^ ions are important factors in the cytotoxicity of PfHRP2; moreover, PfHRP2 cytotoxicity was suppressed by a metal ion chelator, ethylenediaminetetraacetic acid (EDTA), which is an approved and inexpensive drug used to treat metal ion-associated diseases. Collectively, the results of this study show for the first time that PfHRP2 is a pathogenic factor involved in the severity of *P. falciparum* malaria and they provide new insights into the treatment of *P. falciparum* malaria via a method that can neutralize PfHRP2 pathogenicity.

## Results

### Cell Penetration by PfHRP2

Recombinant PfHRP2 was expressed in *Escherichia coli* with a C-terminal FLAG tag (*SI Appendix*, Fig. S1) and purified using anti-FLAG antibody-immobilized affinity resin (Fig. 1*B*). Of note, PfHRP2 was detected not only by anti-PfHRP2 and anti-FLAG antibodies but also by anti-His-tag antibody that binds to six consecutive histidine residues. This finding indicates that PfHRP2 has consecutive histidine sequences in its primary structure. In our previous study, we showed that polyhistidine peptide-fusion proteins efficiently penetrate the cell membrane of the human fibrosarcoma HT1080 cell line (1). Thus, in the present study, we attempted to verify the cell penetration of PfHRP2 in the same HT1080 cells. As expected, flow cytometric analysis showed that fluorescein-labeled PfHRP2 (Fluo-PfHRP2) was internalized into HT1080 cells within a brief period (3 h) (Fig. 1*C*). The efficient cellular uptake of PfHRP2 could be explained by the presence of consecutive histidine sequences detected by the anti-His-tag antibody. In contrast, fluorescein-labeled bovine serum albumin, used as a negative control, showed no cell penetration. A lactic dehydrogenase (LDH) leakage assay showed no LDH leakages from the HT1080 cells treated with PfHRP2 (Fig. 1*D*). Confocal laser scanning microscopy (CLSM) also showed the obvious cellular uptake of Fluo-PfHRP2 in HT1080 cells (Fig. 1*E*). Thus, we demonstrated that PfHRP2 passes across the cell membrane without causing cellular membrane damage.

### Cell-penetrating Mode of PfHRP2 in Human Fibrosarcoma HT1080 cells

To further investigate the cell-penetrating action of PfHRP2 within the observed 3-h period, the effects of pH, temperature, serum, and protein concentration were examined in Fluo-PfHRP2 cellular uptake in HT1080 cells. Cell penetration of Fluo-PfHRP2 increased when PfHRP2 had a positive charge in weak acidic pH conditions (pH 5.8–6.8) (Fig. 2*A*). This indicates that a positive charge may be an important factor for PfHRP2 cell penetration. When assessing the effects of temperature, the cell penetration of Fluo-PfHRP2 was inhibited at 30°C or 20°C compared with cell penetration during incubation at 37°C (Fig. 2*B*), suggesting that PfHRP2 cell penetration depends on a temperature-sensitive cellular uptake pathway. Conversely, the cell penetration of Fluo-PfHRP2 was significantly increased with lower concentrations of fetal bovine serum (FBS) (Fig. 2*C*); thus, PfHRP2 may have been trapped by serum proteins in the presence of FBS. An ultrafiltration assay showed that Fluo-PfHRP2 efficiently bound to FBS components (Fig. 2*D*), indicating that PfHRP2 was weakly trapped by serum in the blood, which in turn affected cell penetration of the protein. Furthermore, the cell penetration of Fluo-PfHRP2 increased in a dose-dependent manner (Fig. 2*E*); however, with higher concentrations of Fluo-PfHRP2 (>3.5 μM), penetration into living cells was not detected in flow cytometric analysis due to an increase in cell death. Thus, we conclude that PfHRP2 is cytotoxic against HT1080 cells at concentrations ≥3.5 μM.

**Fig. 2.**
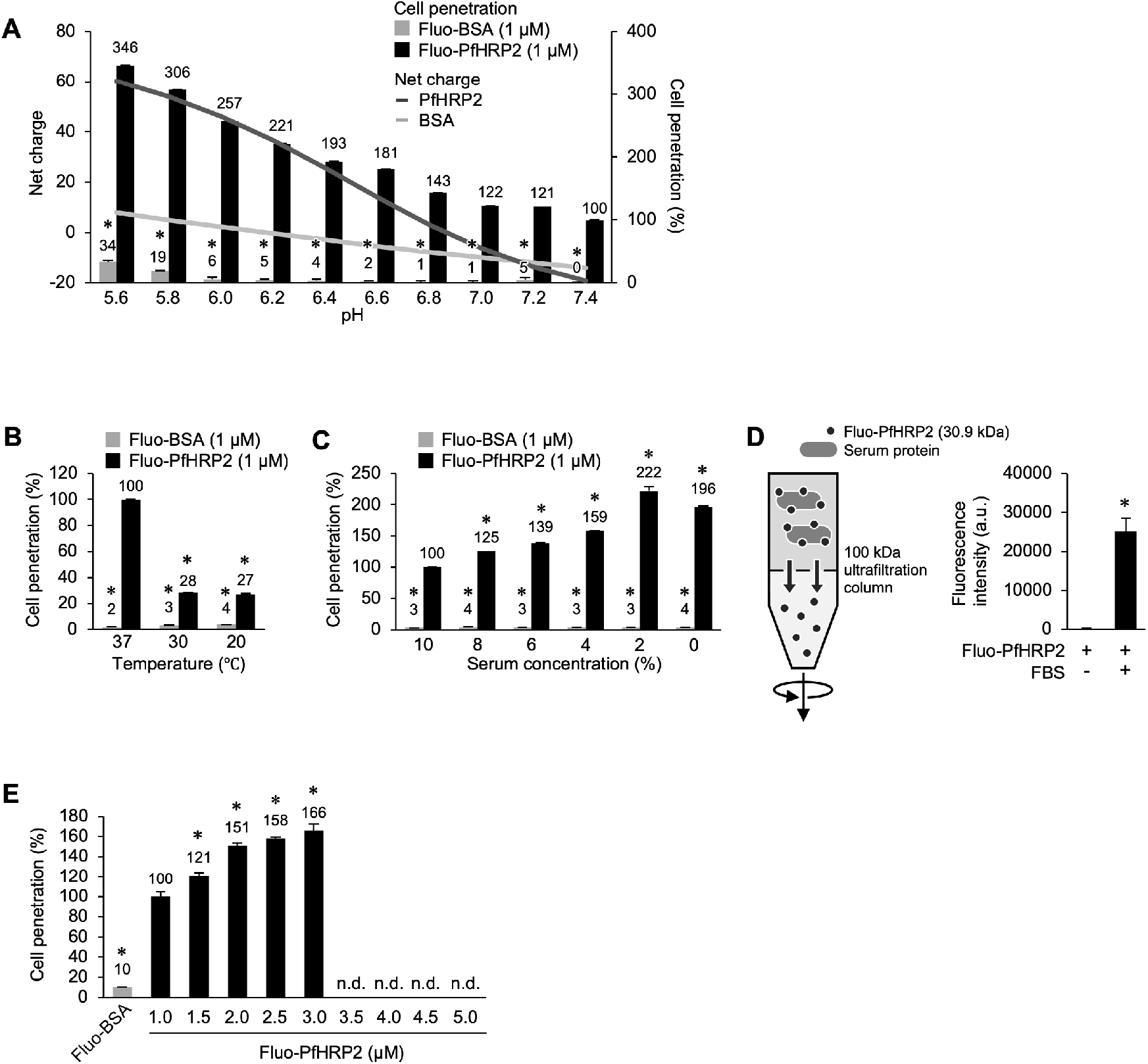
Cellular uptake of PfHRP2 in HT1080 cells. Influences of (*A*) pH, (*B*) temperature, and (*C*) serum concentrations on the cellular uptake of Fluo-PfHRP2 in HT1080 cells. The HT1080 cells were treated with Fluo-PfHRP2 and Fluo-BSA in designed conditions for 3 h. Relative cell penetration of proteins was evaluated by mean fluorescent intensity (MFI) via flow cytometric analysis. (*D*) Binding of Fluo-PfHRP2 and serum proteins. Fluo-PfHRP2 (1 μM) was incubated in 10% fetal bovine serum (FBS) at 37°C for 3 h. Fluo-PfHRP2 (1 μM) bound to serum proteins was collected using a 100-kDa ultrafiltration column. The fluorescence intensity of the ultrafiltrated fraction was quantified using a fluorescence microplate reader. (*E*) Cellular uptake of various concentrations of Fluo-PfHRP2 in HT1080 cells. The HT1080 cells were treated with Fluo-PfHRP2 (1.0–5.0 μM) at 37°C for 3 h. Relative cell penetration was calculated by MFI via flow cytometric analysis. Means and SD are shown (\**p* < 0.05). n.d. indicates “not determined.”

### Cytotoxicity of PfHRP2 against HT1080 cells

To assess the cytotoxicity of PfHRP2 against HT1080 cells, we conducted cell viability assays including PfHRP2 under various conditions. Predictably, PfHRP2 showed dose-dependent cytotoxicity at concentrations up to 3.5 μM over a short period (3 h) (Fig. 3*A*). Thus, we predicted that PfHRP2 may show cytotoxicity at lower concentrations over a longer exposure period. A low concentration of PfHRP2 (i.e., 1.0 μM) led to cell penetration within 3 h; 1.0-μM PfHRP2 was not cytotoxic over a short period (3 h) but showed significant cytotoxicity over longer periods (12 and 24 h) (Fig. 3*B*). Thus, PfHRP2 apparently shows cytotoxicity after cell penetration. Furthermore, the cytotoxicity of PfHRP2 at 1.0 μM for 24 h was greatly increased in the absence of FBS (Fig. 3*C*), which suggests that in the absence of serum conditions, e.g., during malnutrition, the risk of PfHRP2 cytotoxicity would increase. Because PfHRP2 exists at concentrations of 30–100 nM in *P. falciparum* malaria-infected patients’ blood (12–15), we also examined the cytotoxicity of PfHRP2 at these native concentrations for several days. Interestingly, at higher native concentrations of PfHRP2 (90–100 nM), the protein was significantly cytotoxic for 5 days (Fig. 3*D*). Similar PfHRP2 cytotoxicity was also observed over 3 days of incubation (*SI Appendix*, Fig. S2). Therefore, these results show for the first time that PfHRP2 is a slow-acting pathogenic factor produced by *P. falciparum* malaria.

**Fig. 3.**
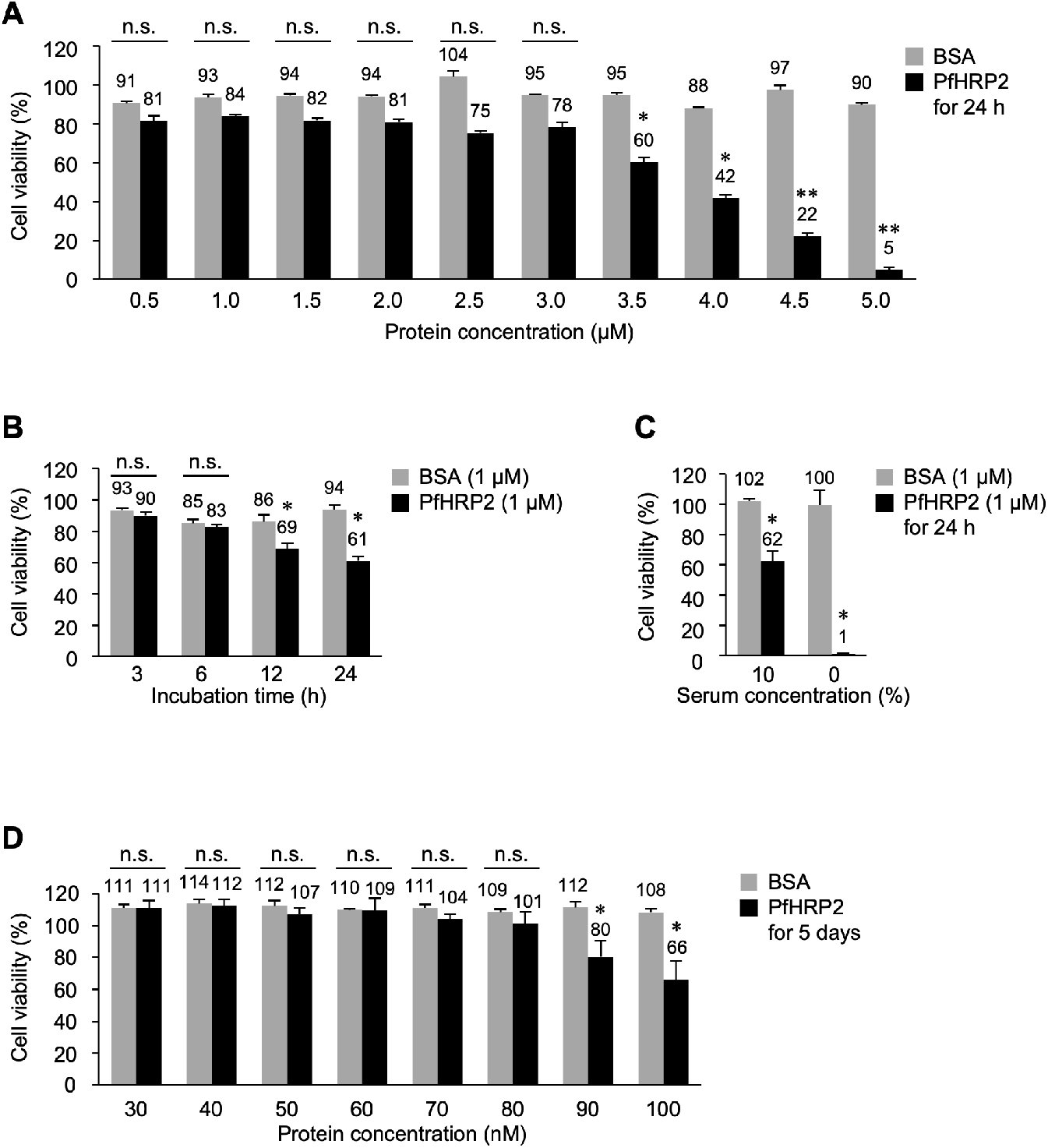
Cytotoxicity of PfHRP2 in HT1080 cells. Influences of (*A*) protein concentration, (*B*) incubation time, and (*C*) serum concentrations on the cytotoxicity of PfHRP2 in HT1080 cells. HT1080 cells were treated with PfHRP2 and BSA in designed conditions at 37°C. (*D*) Cytotoxicity of PfHRP2 at native concentrations for 5 days in HT1080 cells. The HT1080 cells were treated with PfHRP2 at native concentrations (30–100 nM) in *P. falciparum* malaria-infected patients’ blood (maintained at 37°C for 5 days). Relative cell viability was determined using the Cell Counting Kit-8 reaction solution. Means and SD are shown (\**p* < 0.05 and \*\**p* < 0.01). n.s. indicates a nonsignificant difference.

### Cell-penetrating and Cytotoxic Domain of PfHRP2

PfHRP2 showed cell penetration and subsequent cytotoxicity in the experiments described above; thus, we attempted to determine the cell-penetrating and cytotoxic domain of the protein. PfHRP2 mainly consists of the repetitive sequence AHHAHHAAD. First, we designed and synthesized AHHAHHAAD_1_, AHHAHHAAD_2_, AHHAHHAAD_3_, and AHHAHHAAD_4_ peptides consisting of sequences with 1–4 repeats of AHHAHHAAD (*SI Appendix*, Fig. S3). However, these repetitive AHHAHHAAD peptides lacked cytotoxicity in HT1080 cells at concentrations up to 100 μM over 24-h exposures (*SI Appendix*, Fig. S3). Thus, truncated PfHRP2 lacking multiple AHHAHHAAD sequences, i.e., PfHRP2(Δ1–42), (Δ1–108), and (Δ1–204), were prepared via an *E. coli* expression system (Fig. 4*A*) before their cell penetration and cytotoxicity were determined. Compared with the cell penetration of full-length PfHRP2, all truncated PfHRP2 proteins showed reduced cell penetration (50% or less) (Fig. 4*B*). In a cell viability assay, full-length PfHRP2 and PfHRP2(Δ1–42) had similar cytotoxicities, whereas PfHRP2(Δ1–108) and PfHRP2(Δ1–204) lacked cytotoxicity (Fig. 4*C*). These results indicate that the full-length sequence of PfHRP2 is essential for cell penetration with at least the 42–289 amino acid sequence being required for cytotoxicity.

**Fig. 4.**
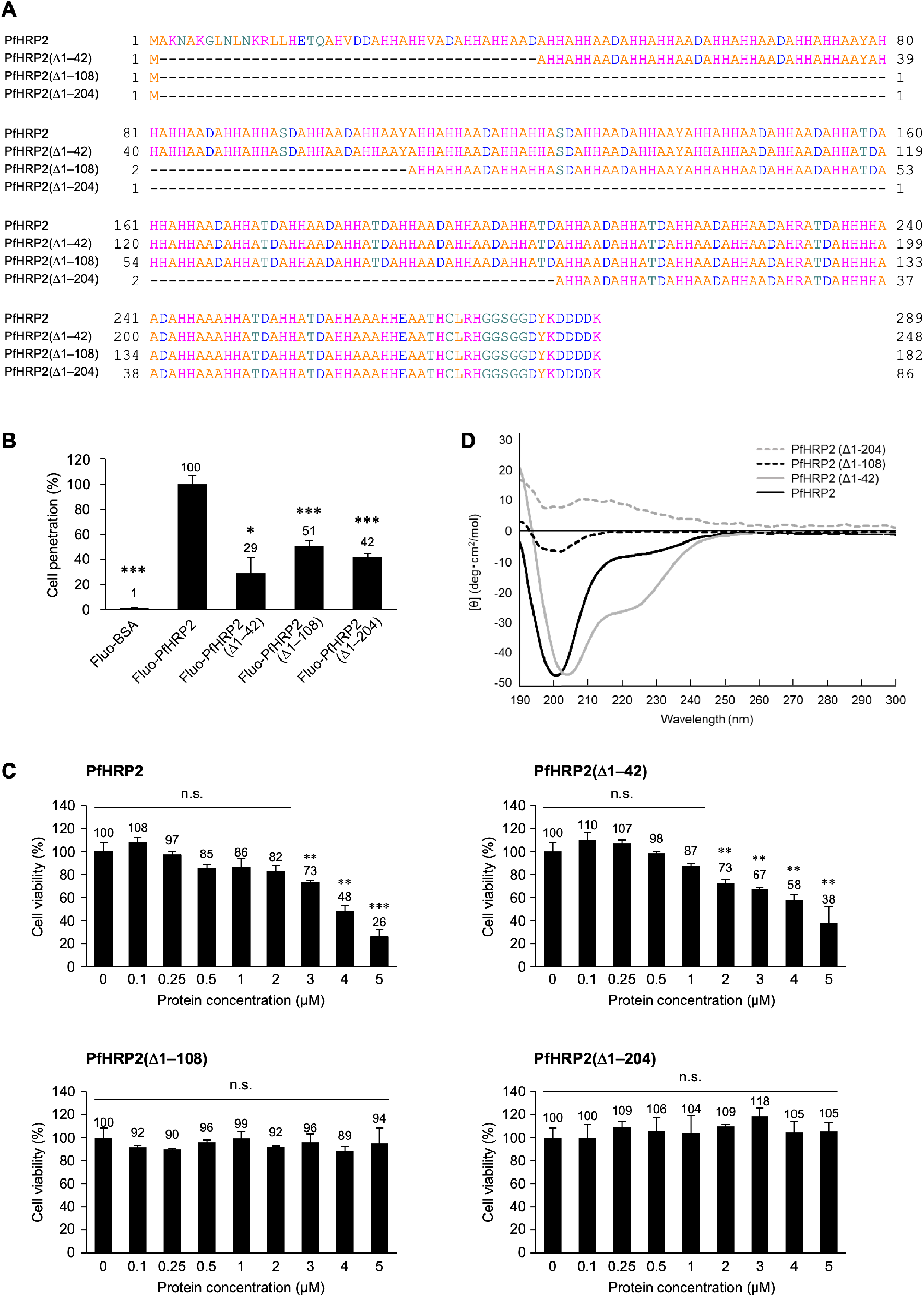
Cell-penetrating and cytotoxic region of PfHRP2. (*A*) Amino acid sequences of full-length and truncated PfHRP2. (*B*) Comparison between cellular uptake of the full-length and truncated PfHRP2. HT1080 cells were treated with the fluorescein-labeled full-length and truncated PfHRP2 at 1 μM and 37°C for 3 h. Relative cell penetration of proteins was calculated by the MFI in the flow cytometric analysis. Means and SD are shown (\**p* < 0.05, \*\**p* < 0.01, and \*\*\**p* < 0.001). (*C*) Comparison between the cytotoxicity of the full-length and truncated PfHRP2. The HT1080 cells were treated with various concentrations of full-length and truncated PfHRP2 at 37°C for 24 h. Relative cell viability was determined using the Cell Counting Kit-8 reaction solution. Means and SD are shown (\**p* < 0.05, \*\**p* < 0.01, and \*\*\**p* < 0.001). (*D*) Circular dichroism (CD) spectra of the full-length and truncated PfHRP2. The CD spectrum of each protein (5 μM) was measured in 20-mM phosphate buffer (pH 7.4) using a 1-mm path length cell at 37°C on a Jasco J-820 CD spectrophotometer.

Circular dichroism (CD) spectrum measurements showed that both full-length PfHRP2 and PfHRP2(Δ1–42), which had the same level of cytotoxicity, had a CD spectrum with a negative peak near 200 nm that is indicative of a random coil secondary structure (Fig. 4*D*). Conversely, the random coil secondary structure was not observed in PfHRP2(Δ1–108) and PfHRP2(Δ1–204), which lacked cytotoxicity. Thus, the random coil secondary structure seems to be vital for the cytotoxicity of PfHRP2. Comparing full-length PfHRP2 with high cell-penetrating ability and PfHRP2(Δ1–42) with moderate cell-penetrating ability, it was observed that the latter had a CD spectrum with a larger negative peak near 222 nm, which is indicative of an α-helix secondary structure (Fig. 4*D*); this difference suggests that the excessive α-helix secondary structure inhibits the cell penetration of PfHRP2.

### Cytotoxic Mechanism of PfHRP2 in HT1080 cells

PfHRP2 showed cytotoxicity without cell membrane damage in the LDH leakage assay (Fig. 1*D*); thus, we eliminated the necrosis pathway as a candidate mechanism of PfHRP2 cytotoxicity. To identify the mechanism, we first investigated the possibility that PfHRP2 kills human cells via apoptosis. However, the human fibrosarcoma HT1080 cells treated with PfHRP2 showed no morphological changes (*SI Appendix*, Fig. S4*A*). An apoptosis detection assay, the Annexin V assay, also indicated that PfHRP2 did not induce phosphatidylserine exposure, which is a known apoptosis marker (*SI Appendix*, Fig. S4*B*). Thus, we examined the possibility that PfHRP2 induces autophagy-associated cytotoxicity. Basal autophagy maintains cell proliferation by degradation of intracellular components and the suppression of basal autophagy has been reported to reduce cell proliferation (16). Our CLSM analysis revealed that PfHRP2 localized to intracellular lysosomes, which are known autophagy-associated organelles, after cell penetration (Fig. 5*A*). Interestingly, PfHRP2 inhibited autolysosome formation, an event that is induced by lysosome–autophagosome fusion and is essential for the subsequent autophagy process, in a similar manner to a typical autophagy inhibitor, bafilomycin A1 (Fig. 5*B*). Furthermore, in cells treated with PfHRP2, the amounts of p62 protein, a metabolite accumulated by autophagy dysfunction, increased in a time-dependent manner (Fig. 5*C*). These results indicate that PfHRP2 is internalized into lysosomes and suppresses basal autophagy by disturbing autolysosome formation (i.e., lysosome–autophagosome fusion) in the cells, similar to bafilomycin A1 (Fig. 5*D*).

**Fig. 5.**
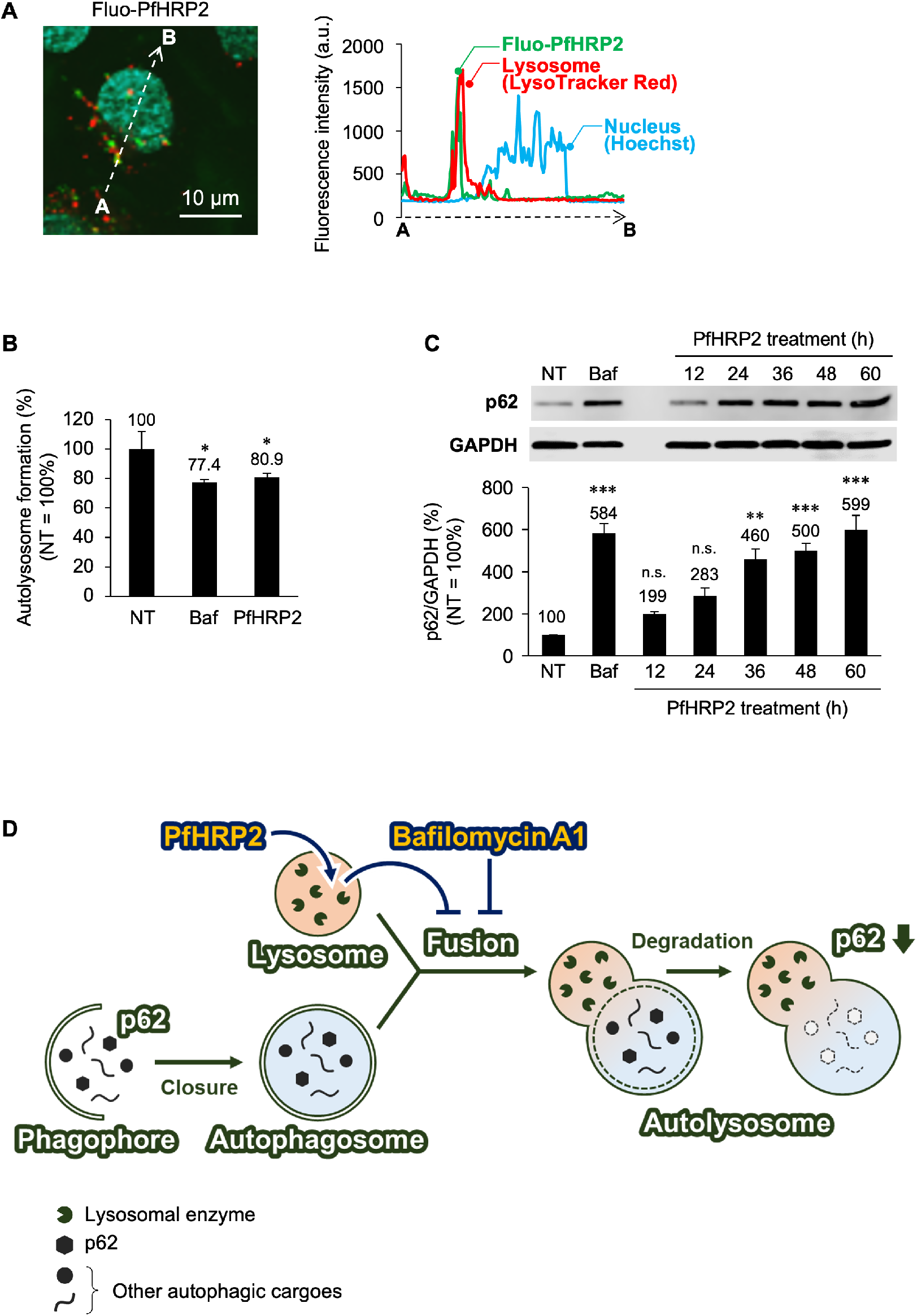
Autolysosome dysfunction by PfHRP2 in HT1080 cells. (*A*) Colocalization of Fluo-PfHRP2 to lysosomes in HT1080 cells. Distribution of the Fluo-PfHRP2, nucleus, and lysosome was quantified by measuring fluorescence intensity in the dotted line areas between A and B. Green, red, and cyan fluorescence indicate Fluo-PfHRP2, endogenous lysosome stained with LysoTracker Red, and nucleus stained with Hoechst 33258, respectively. Scale bar: 10 μm. (*B*) Autolysosome formations inhibited by PfHRP2 in HT1080 cells. The HT1080 cells were treated with bafilomycin A1 (Baf; 25 nM), a typical inhibitor of autolysosome formation, or PfHRP2 (1 μM) at 37°C for 12 h. Cellular autolysosomes were stained by the DAL Green autolysosome-specific fluorescent probe and relative autolysosome formation was calculated by MFI in the flow cytometric analysis. (*C*) Autolysosome dysfunction induced by PfHRP2 in HT1080 cells. The HT1080 cells were treated with bafilomycin A1 (Baf; 100 nM) for 4 h or PfHRP2 (1 μM) for 12–60 h at 37°C, respectively. Cell lysates were prepared from the treated cells, and p62 protein, a metabolite accumulated during autolysosome dysfunction, was detected by western blotting. Relative amounts of p62 protein were calculated by normalizing p62 with glyceraldehyde 3-phosphate dehydrogenase (GAPDH). Means and SD are shown (\**p* < 0.05, \*\**p* < 0.01, and \*\*\**p* < 0.001). n.s. indicates a nonsignificant difference. (*D*) Scheme for PfHRP2 inhibiting cellular proliferation. PfHRP2 is internalized into lysosomes and inhibits autolysosome formation in a similar manner to bafilomycin A1.

### Ca^2+^-dependent PfHRP2 Cytotoxicity and the Protective Effect of EDTA in HT1080 cells

In an attempt to prevent PfHRP2 pathogenicity, we searched for a potential inhibitor against the protein. First, we assessed the repetitive AHHAHHAAD peptides consisting of partial sequences in PfHRP2 to test whether repetitive AHHAHHAAD peptides show competitive inhibition of pathogenic PfHRP2. However, all repetitive AHHAHHAAD peptides lacked inhibitory effects on PfHRP2 cytotoxicity in HT1080 cells (*SI Appendix*, Fig. S3). Anti-PfHRP2 and anti-His tag antibodies also lacked competitive inhibition in terms of PfHRP2 cell penetration and cytotoxicity (*SI Appendix*, Fig. S5). Interestingly, however, the metal ion chelator EDTA and ethylene glycol tetraacetic acid (EGTA) significantly inhibited the cytotoxicity of PfHRP2 (Fig 6*A*). Conversely, neither EDTA nor EGTA inhibited the cell penetration ability of PfHRP2 (*SI Appendix*, Fig. S6). EDTA is known as a chelator for various divalent metal ions whereas EGTA is a Ca^2+^-specific chelator. This result suggests that Ca^2+^ ions are essential for the cytotoxicity of PfHRP2 but not its cell penetration. Thus, the relationship between Ca^2+^ ions and PfHRP2 pathogenicity was determined. A metal ion detection assay showed that Ca^2+^ ions bound to PfHRP2 and their binding was inhibited in the presence of EDTA (Fig. 6*B*). Furthermore, the influence of Ca^2+^ ions on PfHRP2 cytotoxicity was examined in a Ca^2+^ concentration-modified medium (*SI Appendix*, Table S1). Human serum Ca^2+^ concentrations were 78.1–109.0 mg/L (17); thus, we used a modified medium containing up to 100 mg/L of Ca^2+^ ions. As expected, Ca^2+^ ions enhanced PfHRP2 cytotoxicity in a dose-dependent manner (Fig. 6*C*). However, CD spectrum analysis revealed negligible changes in the secondary structure of PfHRP2 in the presence of Ca^2+^ or EDTA (*SI Appendix*, Fig. S7). Therefore, PfHRP2 cytotoxicity can be attributed to factors affected by Ca^2+^ ions rather than to the protein’s secondary structure. As previously mentioned, we showed that PfHRP2 localized to lysosomes (Fig. 5*A*). Furthermore, lysosomal Ca^2+^ homeostasis is crucial for cell proliferation (18); thus, we assessed lysosomal Ca^2+^ levels in cells treated with PfHRP2 and EDTA. As expected, PfHRP2 treatment increased the amount of lysosomal Ca^2+^ in a dose-dependent manner (Fig. 6*D*), whereas the PfHRP2-induced increase in lysosomal Ca^2+^ levels was inhibited by EDTA (Fig. 6*E*). Generally, Ca^2+^ ions released from lysosomes via a Ca^2+^-permeable channel promote autophagy (19). However, a recent study reported that lysosomal Ca^2+^ not only acts as a positive regulator but also functions as a negative regulator in autophagy through the reactivation of mTOR kinase (20). Another study also reported that high Ca^2+^ concentrations in cytosol inhibit autolysosome formation (21). Given that these studies report Ca^2+^ ions as negative regulators of autophagy, our findings suggest that PfHRP2 cytotoxicity arises due to an increase in lysosomal Ca^2+^ (or a subsequent increase in cytosolic Ca^2+^), which acts as a negative regulator of autophagy. Thus, Ca^2+^ seems to be an essential factor for PfHRP2 cytotoxicity and the pathogenicity of PfHRP2 appears to be caused by increasing lysosomal Ca^2+^ levels. Furthermore, our findings suggest that Ca^2+^ ion-chelators (e.g., EDTA and EGTA) are promising candidates as PfHRP2 inhibitors. Based on our results, we propose a model for the pathogenic and protective mechanisms of PfHRP2 and EDTA, respectively, as shown in Fig. 7.

**Fig. 6.**
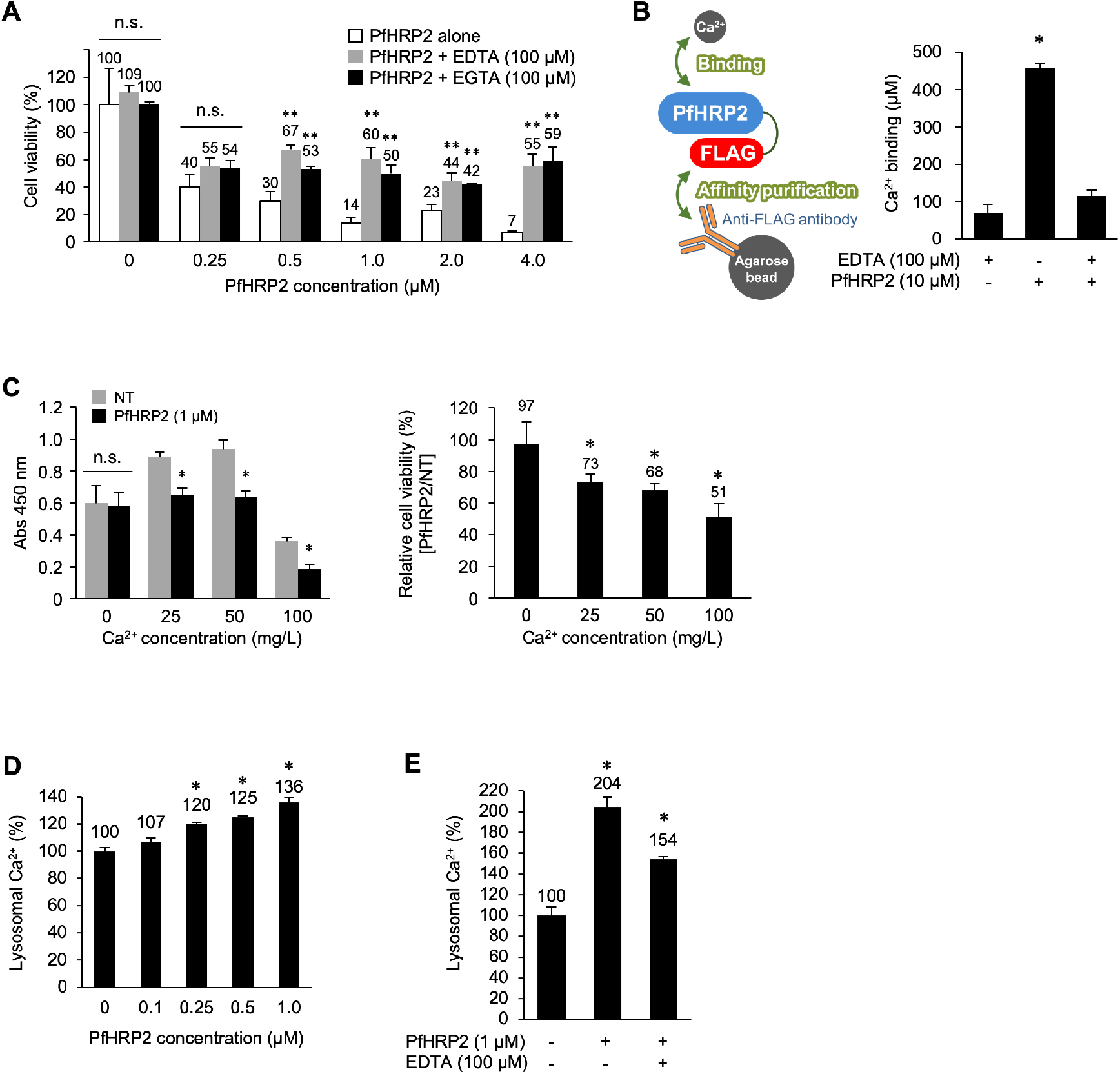
Protective effects of EDTA/EGTA on Ca^2+^-dependent PfHRP2 cytotoxicity. (*A*) Protective effects of EDTA/EGTA on PfHRP2 cytotoxicity in HT1080 cells. HT1080 cells were treated with PfHRP2 (1 μM) in the presence or absence of EDTA/EGTA (100 μM) at 37°C for 24 h. Relative cell viability was determined using the Cell Counting Kit-8 reaction solution. (*B*) Binding of PfHRP2 and Ca^2+^ ions. The PfHRP2 (10 μM) was incubated in the Eagle’s minimal essential medium containing CaCl_2_ (200 mg/L) with/without EDTA (100 μM) at 37°C for 1 h. Ca^2+^ ion bound to the PfHRP2 was collected using anti-FLAG antibody-immobilized resin. Amounts of collected Ca^2+^ ions were determined using a Metallo Assay LS Trace Metal Assay Kit for calcium. (*C*) Ca^2+^-dependent cytotoxicity of PfHRP2 in HT1080 cells. HT1080 cells were treated with PfHRP2 (1 μM) in the modified medium containing various concentrations of Ca^2+^ (0–100 mg/L) at 37°C for 24 h. Relative cell viability was determined using the Cell Counting Kit-8 reaction solution. (*D*) Increase of lysosomal Ca^2+^ by PfHRP2 and (*E*) inhibitory effect of EDTA in HT1080 cells. HT1080 cells were treated with various concentrations of PfHRP2 (0.1–1.0 μM) with or without EDTA (100 μM) at 37°C for 24 h. Lysosomal Ca^2+^ were stained with Cal520-Dextran (MW 10,000), a lysosomal Ca^2+^-specific fluorescent indicator, and then relative lysosomal Ca^2+^ levels were determined by measuring MFI in the flow cytometric analysis. Means and SD are shown (\**p* < 0.05 and \*\**p* < 0.01). n.s. indicates a nonsignificant difference.

**Fig. 7.**
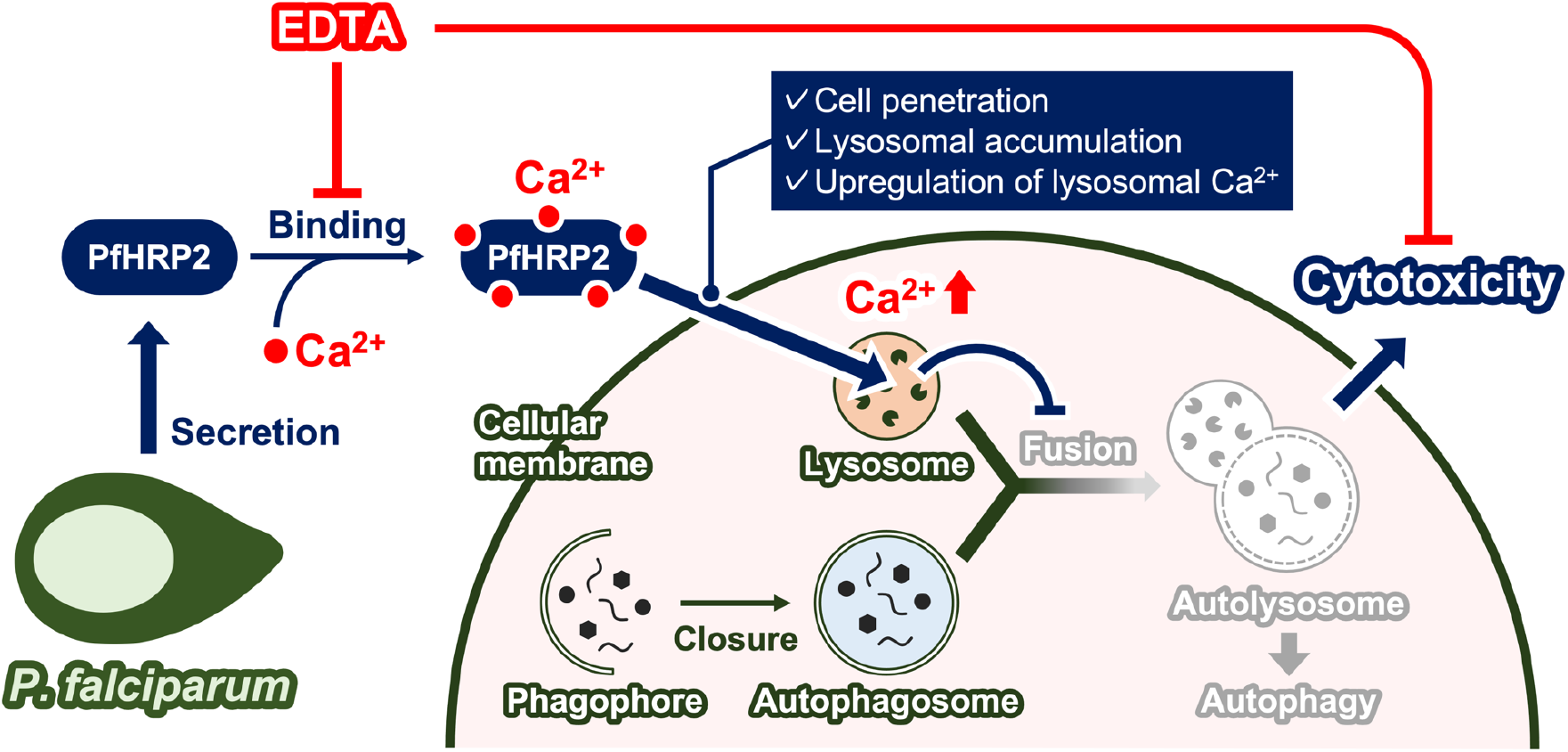
A model for the pathogenic and protective mechanisms of PfHRP2 and EDTA. *Plasmodium falciparum* secretes PfHRP2 to the bloodstream in *P. falciparum*-malaria patients. The secreted PfHRP2 binds to Ca^2+^ ions and penetrates the cell membrane. After cell penetration, the PfHRP2 with Ca^2+^ accumulates to the lysosome, and the lysosomal Ca^2+^ level increases. Lysosomal Ca^2+^ homeostasis is disturbed by PfHRP2 and lysosome–autophagosome fusion is inhibited. Finally, autolysosome formation, essential for the basal autophagy that maintains cellular metabolism, is also suppressed and cellular proliferation is inhibited. However, EDTA inhibits the binding between PfHRP2 and Ca^2+^ ions. Thus, EDTA also inhibits the increase of lysosomal Ca^2+^ levels and subsequent cytotoxicity.

### PfHRP2 Pathogenicity and the Protective Effect of EDTA in Various Human Cells

We demonstrated PfHRP2 pathogenicity and EDTA protective effects in human fibrosarcoma HT1080 cells, but PfHRP2 and EDTA also showed varying degrees of cytotoxicity and protective effects, respectively, in various human cell lines including HT1080 cells (Fig. 8*A*). PfHRP2 reduced cell proliferation by 87% in HepG2 human hepatoma cells, 43% in HT1080 human fibrosarcoma cells, 34% in MKN74 human gastric cancer cells, 29% in Caco-2 human colon carcinoma cells, and 19% in RERF-LC-Al human squamous lung cancer cells. In contrast, EDTA ameliorated PfHRP2 cytotoxicity by 55% in HepG2 human hepatoma cells, 41% in HT1080 human fibrosarcoma cells, and 24% in MKN74 human gastric cancer cells. However, EDTA was ineffective against PfHRP2 cytotoxicity in RERF-LC-Al human squamous lung cancer cells and Caco-2 human colon carcinoma cells. Thus, the human cell lines used in the present study can be divided into two groups: the EDTA-effective group (HT1080, MKN74, and HepG2 cells) and EDTA-ineffective group (RERF-LC-Al and Caco-2 cells).

**Fig. 8.**
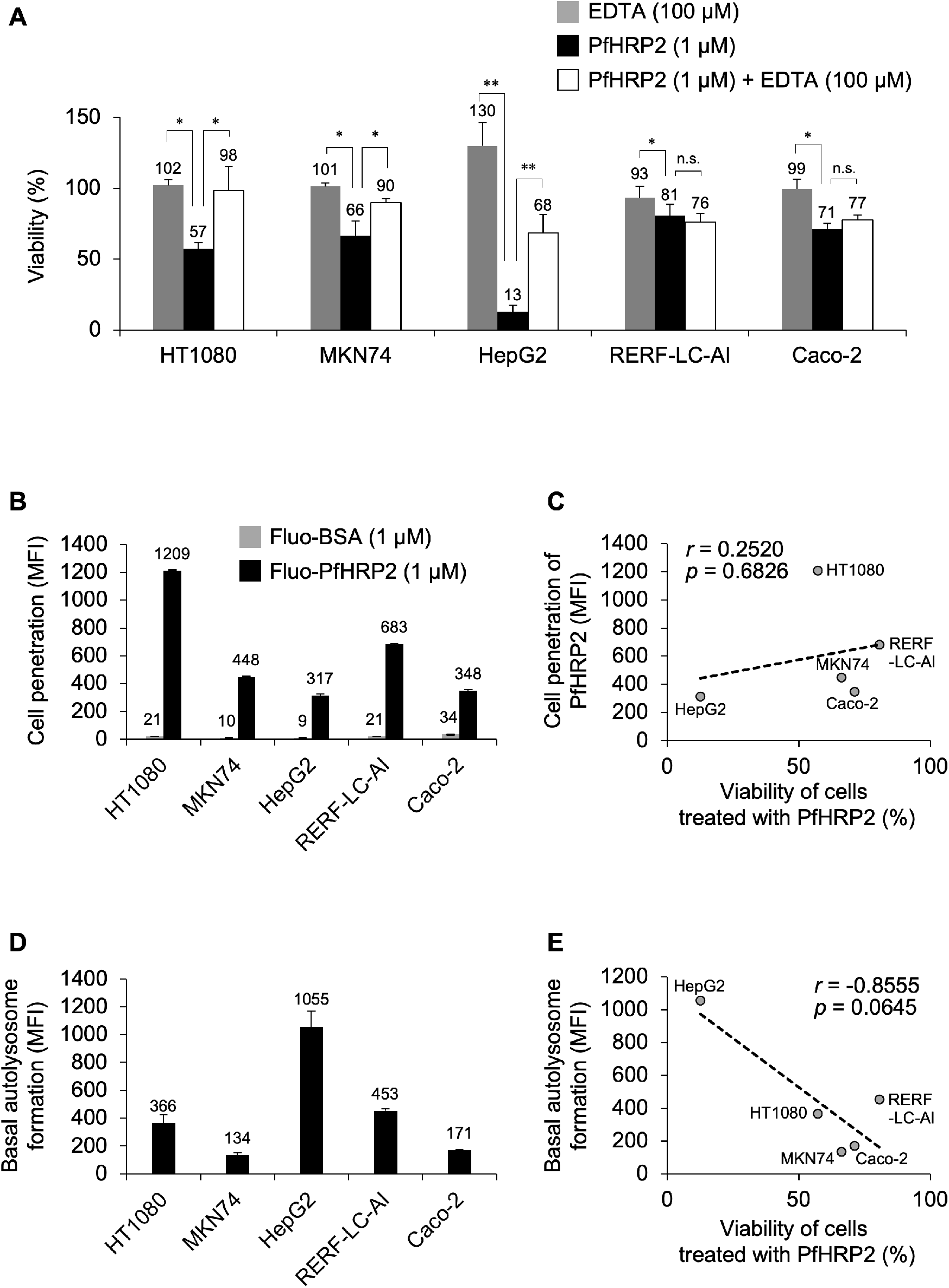
PfHRP2 pathogenicity and the protective effect of EDTA in various human cells. (*A*) PfHRP2 cytotoxicity and protective effect of EDTA in various human cells. Cells were treated with PfHRP2 (1 μM) in the presence or absence of EDTA (100 μM) at 37°C for 24 h. Relative cell viability was determined using the Cell Counting Kit-8 reaction solution. (*B*) Cellular uptake of PfHRP2 in various human cells. Cells were treated with the Fluo-PfHRP2 (1 μM) and Fluo-BSA (1 μM) at 37°C for 3 h. Cell penetration of proteins was evaluated by mean fluorescent intensity (MFI) in the flow cytometric analysis. (*C*) Correlated distribution between PfHRP2 cytotoxicity and cell penetration. The X-axis indicates relative the viability of cells treated with PfHRP2 (%), see Fig. 8*A*, whereas the Y-axis indicates cell penetration of PfHRP2 (MFI), see Fig. 8*B*. r indicates the Pearson’s coefficient of correlation. (*D*) Basal autolysosome formation in various human cells. Cellular autolysosomes were stained with DAL Green and autolysosome formation was determined by MFI in the flow cytometric analysis. (*E*) Correlated distribution between PfHRP2 cytotoxicity and basal autophagy level. The X-axis is the same as in *C*, whereas the Y-axis indicates basal autolysosome formation, see Fig. 8*D*. r indicates the Pearson’s correlation coefficient. Means and SD are shown (\**p* < 0.05 and \*\**p* < 0.01). n.s. indicates a nonsignificant difference. Cell lines: HT1080, human fibrosarcoma cells; MKN74, human gastric cancer cells; HepG2, human hepatoma cells; RERF-LC-Al, human squamous lung cancer cells; and Caco-2, human colon carcinoma cells.

To understand the underlying reason for differences in PfHRP2 cytotoxicity among human cell lines, we determined and compared PfHRP2 cell penetration and basal autophagy levels in the various human cell lines. PfHRP2 showed varying degrees of cell penetration in these cell lines, but the largest amount of cellular uptake was in HT1080 human fibrosarcoma cells (Fig. 8*B*). Pearson’s correlation analysis showed a weak positive correlation (Pearson’s correlation coefficient: *r* = 0.2520) between the cytotoxicity and cell penetration of PfHRP2, but this correlation was not statistically significant (*p* = 0.6826) (Fig. 8*C*). In contrast, a basal autophagy detection assay, the DAL green assay in which autolysosome formation is measured, showed differing levels of basal autophagy in human cell lines (Fig. 8*D*), with the HepG2 human hepatoma cells showing the highest levels of basal autophagy. Interestingly, a strong negative correlation (Pearson’s correlation coefficient: *r* = −0.8555) existed between the viability of cells treated with PfHRP2 and the basal autophagy levels in seven human cell lines (Fig. 8*E*). Although statistical analysis showed that the correlation between the cytotoxicity of PfHRP2 and basal autophagy was not significant, the strong negative trend was close to statistical significance (*p* = 0.0645). These results suggest that PfHRP2 cytotoxicity moderately depends on basal autophagy levels in cells.

## Discussion

Histidine-rich sequences increase the cell penetration of proteins into human cells. Previously, we have demonstrated that nonmembrane permeable proteins, e.g., GFP and GLA, are internalized into human cells by artificial fusion of a polyhistidine peptide consisting of only 16 histidine residues (1, 2). Hence, we hypothesized that natural histidine-rich peptides and proteins may also have similar cell membrane permeability to human cells. In the present study, we demonstrated that PfHRP2, a natural histidine-rich protein produced by *P. falciparum*, showed cell penetration and subsequent cytotoxicity in various human cells. Indeed, this is the first report to demonstrate that a natural histidine-rich protein has cell-penetrating abilities and intracellular function in human cells. As PfRFP2 has such properties, it is possible that other natural histidine-rich proteins may also be internalized into human cells to produce intracellular effects. In future research, further assessment of cell penetration properties may lead to the discovery of new functions for natural histidine-rich proteins other than PfHRP2. Other natural histidine-rich proteins of interest include *Homo sapiens* histidine-rich glycoprotein (HsHRGP). HsHRGP is a secretory protein found in large amounts in human blood (21) and is frequently in contact with human cells. In addition, HsHRGP is known to have multiple functions including involvement in immunity, angiogenesis, coagulation, clearance of apoptotic phagocytes, cell adhesion, migration, angiogenesis, and sepsis (22–25). Hence, new functions of the natural histidine-rich protein related to human health care are expected to be identified by investigating their cell membrane permeation. Although the biomimetic approach has recently become popular for the creation of artificial functional peptides based on natural protein structures (26–30), the present study pioneered a novel “reverse biomimetic approach” to reveal the function of natural proteins based on artificial peptide structures.

Here, we also revealed that the cytotoxicity of PfHRP2 arises by inhibiting basal autophagy after cell penetration. In other words, PfHRP2 is a pathogenic factor produced by *P. falciparum*. In particular, PfHRP2 showed significant cytotoxicity over 3–5 days at concentrations typically found in *P. falciparum*-infected patient’s blood (90–100 nM) (12). This result is consistent with clinical deterioration caused by *P. falciparum* malaria occurring 3–7 days after the onset of fever (6). In addition, the cell penetration and cytotoxicity of PfHRP2 increased 2.5- and 2.6-fold in the absence of serum, respectively. Therefore, low serum protein concentrations, which occur during malnutrition, seem to increase the risk of negative effects from PfHRP2; this finding is consistent with malnutrition increasing the lethality of malaria infection (31–34). We also showed that PfHRP2 was highly cytotoxic to human liver cancer-derived HepG2 cells. The basal level of autolysosome activity in liver cells is known to be higher than that in other tissue cells due to active detoxification and degradation (35), which is consistent with the results of our study. Therefore, we infer that HepG2 cells showed higher susceptibility to PfHRP2 pathogenicity and cytotoxicity because of the inhibition of autolysosome formation. Indeed, hepatic dysfunction is typically severe in *P. falciparum*-infected patients (36), which strongly suggests that PfHRP2 plays an important role in the severity of *P. falciparum* malaria. Until now, PfHRP2 has been considered only as a diagnostic marker of *P. falciparum* malaria infection because it is released in high quantities into the patient’s bloodstream. Recently, however, it was reported that PfHRP2 perturbs the blood–brain barrier and promotes the movement of *P. falciparum* into the brain (37, 38). Thus, PfHRP2 is not only a secretory protein but also a harmful protein produced by *P. falciparum*. Taken together, the studies to date suggest that PfHRP2 plays a major role in the severity of *P. falciparum* malaria and may be an important target molecule for the treatment of this disease.

In the present study, we also attempted to elucidate the cytotoxicity mechanism of PfHRP2. We found that it binds to Ca^2+^ ions and that Ca^2+^ ions are required for its cytotoxicity. The AHHAHHAAD peptide, a partial sequence of PfHRP2, has previously been identified as a metal-binding peptide (39); thus, Ca^2+^ is considered an important factor for PfHRP2. Furthermore, the cytotoxicity of PfHRP2 was robust and not entirely neutralized by partial peptides or antibodies. However, EDTA, a common metal ion chelator, effectively suppressed the cytotoxicity of PfHRP2 in some human cell lines. EDTA is already approved as an inexpensive drug for metal ion-associated diseases (40–42); hence, it is expected to be used as a neutralizing agent against PfHRP2 cytotoxicity. In 2019, 94.0% of global malaria deaths occurred in African regions without medical resources and 99.7% of these cases were attributed to *P. falciparum* malaria (7). Therefore, EDTA, as an inexpensive and approved drug, has the potential for use as a treatment to neutralize PfHRP2 cytotoxicity and minimize the severity of *P. falciparum* malaria. Future *in vivo* studies will be required to demonstrate the neutralizing effect of EDTA on PfHRP2 cytotoxicity.

We found that EDTA was unable to completely suppress PfHRP2 cytotoxicity because of its inability to inhibit the cell penetration of PfHRP2. Unfortunately, in the present study, neither antibodies nor repetitive AHHAHHAAD peptides were able to inhibit the cytotoxicity of PfHRP2. To completely suppress the cytotoxicity of PfHRP2, it will be necessary to identify binding partners against PfHRP2 on the cell membrane surface. If such a binding partner is identified, it may be possible to develop an inhibitor that completely suppresses the cell penetration of PfHRP2 by inhibiting the binding between PfHRP2 and its binding partner on the cell membrane surface.

*P. falciparum* malaria is known to produce histidine-rich proteins other than PfHRP2, namely PfHRP1 and PfHRP3. PfHRP1 is a knob-associated histidine-rich protein that forms knob-like protrusions on the surface membrane of *P. falciparum*-infected erythrocytes (43). However, PfHRP1 has a lower histidine content (654 amino acid residues with 49 histidine amino acids; 7.5% of the total amino acids) than that of PfHRP2 and it shows low similarity with PfHRP2. In contrast, PfHRP3 is a secretory protein that has high similarity with PfHRP2, and the mature PfHRP3 protein consists of 233 amino acid residues with 76 histidine amino acids (32.6% of the total amino acids) (44). The structural similarity between PfHRP2 and PfHRP3 leads to cross-reaction of anti-PfHRP2 monoclonal antibodies with PfHRP3. Thus, similar to PfHRP2, PfHRP3 may act as a pathogen. Our findings therefore provide possible novel insights into the potential risk of PfHRP3 regarding *P. falciparum* malaria infection.

Alternatively, elimination of PfHRP2 from *P. falciparum*-infected patient’s blood may also be an effective treatment against *P. falciparum* malaria. Currently, treatment for *P. falciparum* malaria involves the use of artemisinin combination therapy, artemether–lumefantrine, artesunate– amodiaquine, artesunate–mefloquine, and dihydroartemisinin–piperaquine to eliminate *P. falciparum* from the patient’s body (45). However, PfHRP2 is reported to be retained in the whole blood of artesunate-treated patients with malaria at higher levels, even post-parasite clearance (12, 46). Our findings indicate that the large amounts of PfHRP2 present in the patient’s body are also an equally important target for elimination. Even if *P. falciparum* is eliminated from the patient’s body, the persistence of pathogenic PfHRP2 may subsequently induce severe disease. However, if PfHRP2 is efficiently eliminated from the patient’s body at the same time as *P. falciparum*, it may be possible to prevent the severe disease of *P. falciparum* malaria.

## Materials and methods

### Expression and purification of PfHRP2

A recombinant protein expression vector, pET-24b(+) (Merck, Darmstadt, Germany), was used to express fusion proteins combining PfHRP2 with C-terminal FLAG tag via *E. coli* protein expression systems (*SI Appendix*, Fig. S1). The plasmid was transformed into the *E. coli* BL21(DE3) strain by a heat-shock procedure. A single transformant colony was inoculated into Luria-Bertani–Kanamycin (100 μg/mL) medium and grown overnight at 37°C. The culture was inoculated at 2–5 mL into fresh medium and grown at 37°C until an absorbance of 0.6 was noted at 600 nm (mid-logarithmic phase). Isopropyl β-D-1-thiogalactopyranoside was added to the culture medium (final concentration: 0.1 mM) and PfHRP2 expression was induced at 20°C for about 3 h before cell retrieval. The collected *E. coli* cells were homogenized by sonication in 20-mM phosphate buffer (pH 7.4) containing 1-mM phenylmethylsulfonyl fluoride and 200-mM sodium chloride. The homogenate was centrifuged at 12,000 rpm for 20 min. PfHRP2 recombinant protein was purified by anti-FLAG antibody-immobilized affinity gel (Medical & Biological Laboratories Co., Ltd., Nagoya, Japan) and the purified PfHRP2 solution was exchanged into 20-mM sodium phosphate (pH 7.4) by dialysis before being stored at 4°C.

The purified PfHRP2 was detected by western blotting. The PfHRP2 was transferred onto polyvinylidene fluoride (PVDF) membranes and probed using the anti-FLAG polyclonal antibody (rabbit IgG), anti-PfHRP2 polyclonal antibody (mouse IgG), and anti-His-tag polyclonal antibody (rabbit IgG) as the first antibodies at rates of 1:50,000, 1:7,500, and 1:5,000, respectively. The horseradish peroxidase-labeled anti-rabbit IgG antibody and anti-mouse IgG antibody were used as the second antibodies at rates of 1:70,000 and 1:10,000, respectively. A standard western blotting protocol was used.

### Peptide purification

The AHHAHHAAD_1_, AHHAHHAAD_2_, AHHAHHAAD_3_, and AHHAHHAAD_4_ peptides were synthesized by solid-phase peptide synthesis and purchased from Biologica (Shanghai, China). All peptides were purified to homogeneity by high-performance liquid chromatography (HPLC) using a Waters 2489 UV/visible detector and 1524 binary pump (Waters, Milford, MA, USA) and a reversed-phase COSMOSIL 5C18-MS-II column (10 mm × 250 mm) (Nacalai Tesque, Kyoto, Japan). The column was run for 50 min at 3 mL/min with a linear gradient from 10% to 60% (v/v) acetonitrile in water containing 0.1% (v/v) trifluoroacetic acid. The purity of peptides was calculated from peak areas of the HPLC chart, and the final purity of the peptides was >95%. The molecular masses were analyzed using matrix-assisted laser desorption/ionization time-of-flight mass spectrometry (Auto FlexII Bruker Daltonics, Billerica, USA).

### Cell culture

HT1080 human fibrosarcoma cells, HepG2 human hepatoma cells, and RERF-LC-Al human squamous lung cancer cells were cultured in Eagle’s minimal essential medium. Conversely, MKN74 human gastric cancer cells were cultured in RPMI1640 medium. All media contained 10% FBS (v/v), 100-μg/mL streptomycin, 100-units/mL penicillin, and 250-ng/mL amphotericin B. Caco-2 human colon carcinoma cells were cultured in Eagle’s minimal essential medium containing 20% FBS (v/v), 100-μg/mL streptomycin, 100-units/mL penicillin, and 250-ng/mL amphotericin B. All human cell lines were purchased from the RIKEN BioResource Research Center (Ibaraki, Japan); they were maintained at 37°C in a humidified 5% CO_2_ incubator and a subculture was performed every 3–4 days.

### Labeling of proteins with fluorescein

To observe the cellular uptake of PfHRP2, PfHRP2, and Bovine serum albumin (BSA) were labeled using the fluorescein 5-maleimide (Tokyo Chemical Industry Co., Ltd., Tokyo, Japan). Proteins (50–100 μM) in labeling buffer (20-mM sodium phosphate, 150-mM NaCl, and 10-mM EDTA at pH 7.2) were mixed with 25-fold amounts of fluorescein 5-maleimide. The mixtures were shaded and incubated overnight at room temperature. After incubation, fluorescein-labeled proteins were replaced with 20-mM sodium phosphate (pH 7.2) by dialysis.

### LDH leakage assay

To determine cellular membrane damage, cultured cells were seeded onto 96-well plates at a density of 1.0 × 10^4^ cells/well to a final volume of 100 μL and then incubated for 24 h at 37°C in 5% CO_2_. After complete adhesion, the culture media was replaced with media containing 1 μM of proteins and incubated for 3 h. The culture media of each well was replaced with fresh media containing Cytotoxicity LDH Assay Kit-WST (Dojindo, Kumamoto, Japan). LDH leakages, as an index of cellular membrane damage, were determined according to the manufacturer’s guidelines. The absorbance at 490 nm was measured using a multi-plate reader (Tecan, Mannedorf, Switzerland).

### Cell viability assay

For the cellular cytotoxicity assay, cultured cells were seeded onto 96-well plates at a density of 1.0 × 10^4^ cells/well to a final volume of 100 μL and then incubated for 24 h at 37°C in 5% CO_2_. After complete adhesion, the culture media was replaced with fresh media containing various concentrations of proteins and incubated for designed periods. After treatment, the culture media of each well was replaced with 100 μL of fresh media containing the Cell Counting Kit-8 reaction solution (Dojindo) and incubated for 4 h. The absorbance at 450 nm was measured using the multi-plate reader described above.

### Flow cytometric analysis

Cell penetration of fluorescein-labeled proteins was determined by flow cytometry. The cultured human cells were seeded onto 24-well plates at a density of 1.0 × 10^5^ cells/well to a final volume of 500 μL and then incubated for 24 h at 37°C in 5% CO_2_. The culture media was then replaced with fresh media containing various concentrations of fluorescein-labeled proteins and incubated for designed periods. Cells were washed three times with phosphate-buffered saline (PBS), detached with 0.05% trypsin/0.53-mM EDTA, and suspended in 500 μL of FACS buffer (PBS pH 7.4 including 2% FBS). The fluorescence intensity of fluorescein-labeled proteins internalized into cells was measured by flow cytometric analysis with a BD FACSantoII instrument (Becton, Dickinson and Company, NJ, USA). Approximately 10^4^ cells were collected per specimen with three replicates.

### CLSM observations

Cultured cells were seeded onto a multiwell glass-bottom dish (Matsunami Ind., Osaka, Japan) at a density of 1.0 × 10^4^ cells/well to a final volume of 100 μL and incubated for 24 h at 37°C in 5% CO_2_. After complete adhesion, the culture media was replaced with media containing 1 μM of fluorescein-labeled proteins and incubated for 3 h. The cell penetration of fluorescein-labeled proteins was observed (excitation wavelength: 480 nm; emission wavelength: 530 nm) and fluorescence images were obtained using a confocal laser scanning microscope (FluoView; Olympus, Tokyo, Japan).

Intracellular localization of Fluo-PfHRP2 was determined using organelle markers. LysoTracker Red (Life Technologies, CA, USA) and Hoechst33342 (Dojindo) were used as organelle markers for lysosomes and nuclei, respectively. HT1080 cells were precultured in a multiwell glass-bottom dish under the same conditions described above and then incubated with 1-μM Fluo-PfHRP2 for 3 h at 37°C in 5% CO_2_. Subsequently, 500 nM of LysoTracker Red was added to the cells, which were then incubated for 2 h. Hoechst33342 nuclear staining dye was also added to cells for 1 h. Cells were washed three times with PBS and the intracellular localization of Fluo-PfHRP2 was determined using FluoView.

### Autolysosome detection

Autolysosome formation was determined using the autolysosome-specific fluorescent probe, DAL Green (Dojindo), by flow cytometry. The cultured human cells were seeded onto 24-well plates at a density of 1.0 × 10^5^ cells/well to a final volume of 500 μL and incubated for 24 h at 37°C in 5% CO_2_. The culture media was then replaced with fresh media containing DAL green and incubated for 24 h. Cells were washed three times with PBS, detached with 0.05% trypsin/0.53-mM EDTA, and suspended in 500 μL of FACS buffer (PBS pH 7.4 including 2% FBS). The fluorescence intensity of autolysosomes stained by DAL Green was measured by flow cytometric analysis with a BD FACSantoII instrument (Becton, Dickinson and Company). Approximately 10^4^ cells were collected per specimen with three replicates.

### Autophagy detection

The p62 protein, a classical receptor of autophagy and autophagic substrate, was used as a marker to detect autophagy. The HT1080 cells were seeded onto 96-well plates at a density of 1.0 × 10^4^ cells/well to a final volume of 100 μL and then incubated for 24 h at 37°C in 5% CO_2_. After complete adhesion, the culture media was replaced with media containing PfHRP2 (1 μM) and incubated for designed periods. After treatment, the HT1080 cells were lysed in the lysis buffer (20-mM sodium phosphate, pH 7.2; 150-mM sodium chloride; 1% Nonidet P-40; 0.5% sodium deoxycholate; and 0.1% SDS) and the p62 protein was detected by western blotting. The lysates were run on an SDS-PAGE, transferred onto a PVDF membrane, and probed using anti-p62 polyclonal antibody (rabbit IgG) and HRP-labeled anti-rabbit IgG antibody at ratios of 1:5,000 and 1:10,000, respectively. Glyceraldehyde 3-phosphate dehydrogenase (GAPDH) was also detected as an internal control using HRP-labeled anti-GAPDH polyclonal antibody (mouse IgG) at 1:10,000. A standard western blot protocol was used. Autophagy inhibitory effects were calculated as the ratio of p62 to GAPDH protein levels. Bafilomycin A1 was used as an autophagy inhibitor.

### Ca^2+^ ions binding assay

The binding between PfHRP2 and Ca^2+^ ions was determined by calorimetric analysis. PfHRP2 (1 μM) was incubated in Eagle’s minimal essential medium containing 200 mg/L of Ca^2+^ ions at 37°C for 1 h. PfHRP2 was then purified by anti-FLAG antibody-immobilized affinity gel (Medical & Biological Laboratories Co., Ltd); the amounts of Ca^2+^ ions bound to PfHRP2 were determined using a Metallo Assay LS Trace Metal Assay Kit for calcium (CPZ III; MG Metallogenics, Chiba, Japan) according to the manufacturer’s guidelines. The absorbance at 690 nm was measured using the previously mentioned multi-plate reader.

### Lysosomal Ca^2+^ measurement

Lysosomal Ca^2+^ levels were determined by flow cytometry using the Cal520-Dextran (MW 10,000) (AAT Bioquest, CA, USA), a lysosomal Ca^2+^-specific fluorescent indicator. HT1080 cells were seeded onto 96-well plates at a density of 1.0 × 10^4^ cells/well to a final volume of 100 μL and incubated for 24 h at 37°C in 5% CO_2_. After complete adhesion, the culture media was replaced with media containing PfHRP2 (0.1–1.0 μM) with or without EDTA (100 μM) and incubated for 24 h at 37°C in 5.0% CO_2_. After treatment, cells were washed with Hanks’ Balanced Salt solution (HBSS) (Sigma-Aldrich, MO, USA) and incubated in HBSS containing 1.0 μM of Cal520-Dextran (MW 10,000) for 2 h at 37°C in 5.0% CO_2_. Cells were then washed three times with HBSS, detached with 0.05% trypsin/0.53-mM EDTA, and suspended in 500 μL of FACS buffer (PBS pH 7.4 including 2% FBS). The fluorescence intensity of lysosomal Ca^2+^ stained by Cal520-Dextran (MW 10,000) was measured via flow cytometric analysis with a BD FACSantoII instrument (Becton, Dickinson and Company). Approximately 10^4^ cells were collected per specimen with three replicates.

### CD measurement

CD spectra were recorded in the far UV range, from 300 to 190 nm, on a Jasco J-820 CD spectrophotometer with an attached thermal regulator. CD spectra were acquired for recombinant PfHRP2 proteins (1 μM) in the buffer (20 mM sodium phosphate, pH 7.4) using a 1-mm path length cell at 37°C. The typical parameters used in recording the spectra were as follows: bandwidth: 2 nm; response time: 0.5 s; and scan speed: 50 nm/min. Thirty-two scans were collected, the signal of the buffer was subtracted, and the resulting intensities in millidegrees of the proteins were converted to mean residue molar ellipticity (degrees cm^2^ mol^−1^).

### Statistical analysis

Correlations were determined using Pearson’s correlation analysis and data were fit with a linear regression. Differences between the two groups were analyzed by Student’s *t*-test with a two-tailed distribution, whereas differences among multiple groups were evaluated by Tukey’s test. All statistical analyses were conducted with add-on software for Excel (Excel Statistics 2010; SSRI, Tokyo, Japan). *p* values <0.05 were considered statistically significant.

### Data Availability

All study data are included in the article and *SI Appendix*.

## Acknowledgments

This work was supported in part by a grant from the GSK Japan Research Grant 2017 (to T.I.), the Naito Foundation (to T.I.), and JSPS KAKENHI Grant Number JP17K07771 (to T.I.).

## References

1. T. Iwasaki, et al.,Cellular uptake and in vivo distribution of polyhistidine peptides. J. Control. Release 210, 115–124 (2015).

2. T. Iwasaki, N. Murakami, T. Kawano, A polylysine–polyhistidine fusion peptide for lysosome-targeted protein delivery. Biochem. Biophys. Res. Commun. 533, 905–912 (2020).

3. L. J. Panton, et al.,Purification and partial characterization of an unusual protein of Plasmodium falciparum: histidine-rich protein II. Mol. Biochem. Parasitol. 35, 149–160 (1989).

4. M. Rodriguez-del Valle, et al.,Detection of antigens and antibodies in the urine of humans with Plasmodium falciparum malaria. J. Clin. Microbiol. 29, 1236–1242 (1991).

5. M. E. Sinka, et al.,The dominant Anopheles vectors of human malaria in Africa, Europe and the Middle East: Occurrence data, distribution maps and bionomic précis. Parasites and Vectors 3, 117 (2010).

6. A. Trampuz, M. Jereb, I. Muzlovic, R. M. Prabhu, Clinical review: Severe malaria. Crit. Care 7, 315–323 (2003).

7. World malaria report 2020 (June 8, 2021).

8. M. Al-Awadhi, S. Ahmad, J. Iqbal, Current status and the epidemiology of malaria in the middle east region and beyond. Microorganisms 9, 1–20 (2021).

9. M. E. Parra, C. B. Evans, D. W. Taylor, Identification of Plasmodium falciparum histidine-rich protein 2 in the plasma of humans with malaria. J. Clin. Microbiol. 29, 1629–1634 (1991).

10. V. Desakorn, et al.,Semi-quantitative measurement of Plasmodium falciparum antigen PfHRP2 in blood and plasma. Trans. R. Soc. Trop. Med. Hyg. 91, 479–483 (1997).

11. K. E. Poti, D. J. Sullivan, A. M. Dondorp, C. J. Woodrow, HRP2: Transforming Malaria Diagnosis, but with Caveats. Trends Parasitol. 36, 112–126 (2020).

12. P. A. Ndour, et al.,Measuring the Plasmodium falciparum HRP2 protein in blood from artesunate-treated malaria patients predicts post-artesunate delayed hemolysis. Sci. Transl. Med. 9 (2017).

13. C. M. Kifude, et al.,Enzyme-linked immunosorbent assay for detection of Plasmodium falciparum histidine-rich protein 2 in blood, plasma, and serum. Clin. Vaccine Immunol. 15, 1012–1018 (2008).

14. G. S. Park, et al.,Plasmodium falciparum histidine-rich protein-2 plasma concentrations are higher in retinopathy-negative cerebral malaria than in severe malarial anemia. Open Forum Infect. Dis. 4 (2017).

15. S. Uyoga, et al.,Plasma Plasmodium falciparum Histidine-rich protein 2 concentrations in children with malaria infections of differing severity in Kilifi, Kenya. Clin. Infect. Dis. (2020) https://doi.org/10.1093/cid/ciaa1141 (June 8, 2021).

16. V. O. Kaminskyy, T. Piskunova, I. B. Zborovskaya, E. M. Tchevkina, B. Zhivotovsky, Suppression of basal autophagy reduces lung cancer cell proliferation and enhances caspase-dependent and -independent apoptosis by stimulating ROS formation. Autophagy 8, 1032–1044 (2012).

17. N. Ishizaka, et al.,Serum calcium concentration and carotid artery plaque: a population-based study. J. Cardiol. 39, 151–157 (2002).

18. D. A. East, M. Campanella, Ca2+ in quality control: An unresolved riddle critical to autophagy and mitophagy. Autophagy 9, 1710–1719 (2013).

19. D. L. Medina, A. Ballabio, Lysosomal calcium regulates autophagy. Autophagy 11, 970–971 (2015).

20. D. L. Medina, “Lysosomal calcium and autophagy” in International Review of Cell and Molecular Biology, (Elsevier Inc., 2021) https://doi.org/10.1016/bs.ircmb.2021.03.002.

21. C. Mauvezin, P. Nagy, G. Juhász, T. P. Neufeld, Autophagosome-lysosome fusion is independent of V-ATPase-mediated acidification. Nat. Commun. 6, 7007 (2015).

22. I. K. H. Poon, K. K. Patel, D. S. Davis, C. R. Parish, M. D. Hulett, Histidine-rich glycoprotein: The Swiss Army knife of mammalian plasma. Blood 117, 2093–2101 (2011).

23. R. Simantov, et al.,Histidine-rich glycoprotein inhibits the antiangiogenic effect of thrombospondin-1. J. Clin. Invest. 107, 45–52 (2001).

24. M. Blank, Y. Shoenfeld, Histidine-rich glycoprotein modulation of immune/autoimmune, vascular, and coagulation systems. Clin. Rev. Allergy Immunol. 34, 307–312 (2008).

25. H. Wake, Histidine-rich glycoprotein modulates the blood-vascular system in septic condition. Acta Med. Okayama 73, 379–382 (2019).

26. D. T. Haynie, L. Zhang, W. Zhao, J. S. Rudra, Protein-inspired multilayer nanofilms: science, technology and medicine. Nanomedicine Nanotechnology, Biol. Med. 2, 150–157 (2006).

27. Y. Li, X. Liu, X. Dong, L. Zhang, Y. Sun, Biomimetic design of affinity peptide ligand for capsomere of virus-like particle. Langmuir 30, 8500–8508 (2014).

28. W. Du, X. Hu, W. Wei, G. Liang, Intracellular peptide self-assembly: a biomimetic approach for in situ nanodrug preparation. Bioconjug. Chem. 29, 826–837 (2018).

29. R. Surís-Valls, I. K. Voets, Peptidic antifreeze materials: Prospects and challenges. Int. J. Mol. Sci. 20 (2019).

30. Q. Hou, L. Zhang, Biomimetic design of peptide neutralizer of Ebola virus with molecular simulation. Langmuir 36, 1813–1821 (2020).

31. J. S. Edirisinghe, Infections in the malnourished: With special reference to malaria and malnutrition in the tropics. Ann. Trop. Paediatr. 6, 233–237 (1986).

32. E. G. Kasili, Malnutrition and infections as causes of childhood anemia in tropical africa. J. Pediatr. Hematol. Oncol. 12, 375–377 (1990).

33. J. A. Berkley, et al.,HIV infection, malnutrition, and invasive bacterial infection among children with severe malaria. Clin. Infect. Dis. 49, 336–343 (2009).

34. K. D. J. Jones, J. A. Berkley, Severe acute malnutrition and infection. Paediatr. Int. Child Health 34, S1–S29 (2014).

35. T. Ueno, M. Komatsu, Autophagy in the liver: Functions in health and disease. Nat. Rev. Gastroenterol. Hepatol. 14, 170–184 (2017).

36. A. Bhalla, V. Suri, V. Singh, Malarial hepatopathy. J. Postgrad. Med. 52, 315–320 (2006).

37. P. Pal, et al.,Plasmodium falciparum histidine-rich protein II compromises brain endothelial barriers and may promote cerebral malaria pathogenesis. MBio 7 (2016).

38. P. Pal, et al.,Plasmodium falciparum histidine-rich protein II causes vascular leakage and exacerbates experimental cerebral malaria in mice. PLoS One 12 (2017).

39. J. I. B. Janairo, Peptide-Mediated Biomineralization (Springer, 2016) https://doi.org/10.1007/978-981-10-0858-0 (June 11, 2021).

40. P. M. Wax, Current Use of Chelation in American Health Care. J. Med. Toxicol. 9, 303–307 (2013).

41. T. Born, C. N. Kontoghiorghe, A. Spyrou, A. Kolnagou, G. J. Kontoghiorghes, EDTA chelation reappraisal following new clinical trials and regular use in millions of patients: Review of preliminary findings and risk/benefit assessment. Toxicol. Mech. Methods 23, 11–17 (2013).

42. J. A. Drisko, “Chelation Therapy” in Integrative Medicine: Fourth Edition, (Elsevier, 2018), pp. 1004–1015.e3.

43. J. Gruenberg, D. R. Allred, I. W. Sherman, Scanning electron microscope-analysis of the protrusions (knobs) present on the surface of Plasmodium falciparum-infected erythrocytes. J. Cell Biol. 97, 795–802 (1983).

44. E. P. Rock, et al., Comparative analysis of the Plasmodium falciparum histidine-rich proteins HRP-I, HRP-II and HRP-III in malaria parasites of diverse origin. Parasitology 95, 209–227 (1987).

45. W. G. S. Group, Gametocyte carriage in uncomplicated Plasmodium falciparum malaria following treatment with artemisinin combination therapy: a systematic review and meta-analysis of individual patient data. BMC Med. 14, 79 (2016).

46. K. E. Poti, et al.,In vivo compartmental kinetics of Plasmodium falciparum histidine-rich protein II in the blood of humans and in BALB/c mice infected with a transgenic Plasmodium berghei parasite expressing histidine-rich protein II. Malar. J. 18 (2019).

