## Supplementary Information for "Histidine-rich protein 2: a new pathogenic factor of *Plasmodium falciparum* malaria"

**A**

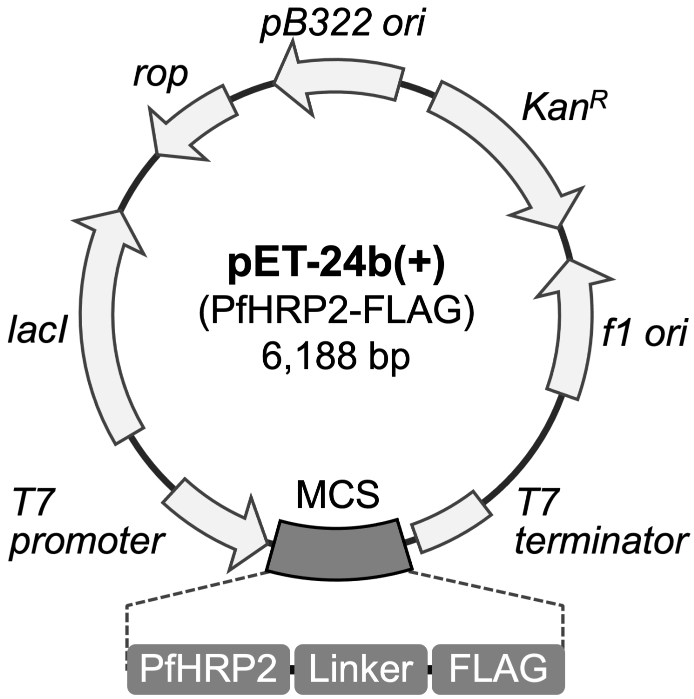


**B**

```
  1 MAKNAKGLNLNKRLLHETQAHVDDAHHAHHVADAHHAHHAADAHHAHHAADAHHAHHAAD
 61 AHHAHHAADAHHAHHAAYAHHAHHAADAHHAHHASDAHHAADAHHAAYAHHAHHAADAHH
121 AHHASDAHHAADAHHAAYAHHAHHAADAHHAADAHHATDAHHAHHAADAHHATDAHHAAD
181 AHHATDAHHAADAHHAADAHHATDAHHAADAHHATDAHHAADAHHAADAHRATDAHHAHH
241 AADAHHAAAHHATDAHHATDAHHAAAHHEAATHCLRHGGSGGDYKDDDDK
```

**Fig. S1.** Recombinant *Plasmodium falciparum* histidine-rich protein 2 (PfHRP2). (*A*) Expression vector for recombinant PfHRP2 including FLAG tag (DYKDDDDK) with linker (GGSGG). The PfHRP2 expression vector was constructed based on the pET-24b(+) vector (Merck, Darmstadt, Germany). The PfHRP2 gene was synthesized to optimize codon usage for *Escherichia coli*. The FLAG tag gene was synthesized by annealing primers with partial homologous sequences and extension. The open circularpET-24b(+) vector was synthesized by inverse polymerase chain reaction (PCR). PCR was performed in the TaKaRa PCR Thermal Cycler Dice Touch (TaKaRa, Shiga, Japan) using the Tks Gflex DNA Polymerase (TaKaRa). The PCR products were ligated into the open circularpET-24b(+) using the In-Fusion HD Cloning Kit (TaKaRa) to generate the PfHRP2 expression vector. The plasmid was sequenced by Eurofins Genomics (Tokyo, Japan) and subsequently transformed into *E. coli* BL21(DE3) (New England BioLabs, MA, USA) for protein expression. (*B*) The amino acid sequence of recombinant PfHRP2 used in the present study. Linker and FLAG-tag are shown as underlining and bold characters, respectively.

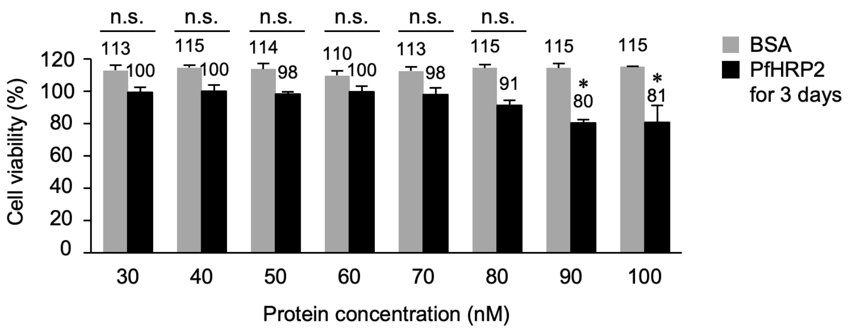


**Fig. S2.** Cytotoxicity of PfHRP2 at native concentrations for 3 days in human fibrosarcoma HT1080 cells. The HT1080 cells were treated with PfHRP2 at a native concentration (30–100 nM) in *P. falciparum* malaria-infected patients' blood (maintained at 37°C for 3 days). Bovine serum albumin (BSA) was used as a negative control. Relative cell viability was determined using the Cell Counting Kit-8 reaction solution. Means and SD are shown (*$p < 0.05$). n.s. indicates a nonsignificant difference.

| Peptide names | Sequences |
| --- | --- |
| (AHHAHHAAD)$_1$ | AHHAHHAAD |
| (AHHAHHAAD)$_2$ | AHHAHHAADAHHAHHAAD |
| (AHHAHHAAD)$_3$ | AHHAHHAADAHHAHHAADAHHAHHAAD |
| (AHHAHHAAD)$_4$ | AHHAHHAADAHHAHHAADAHHAHHAADAHHAHHAAD |

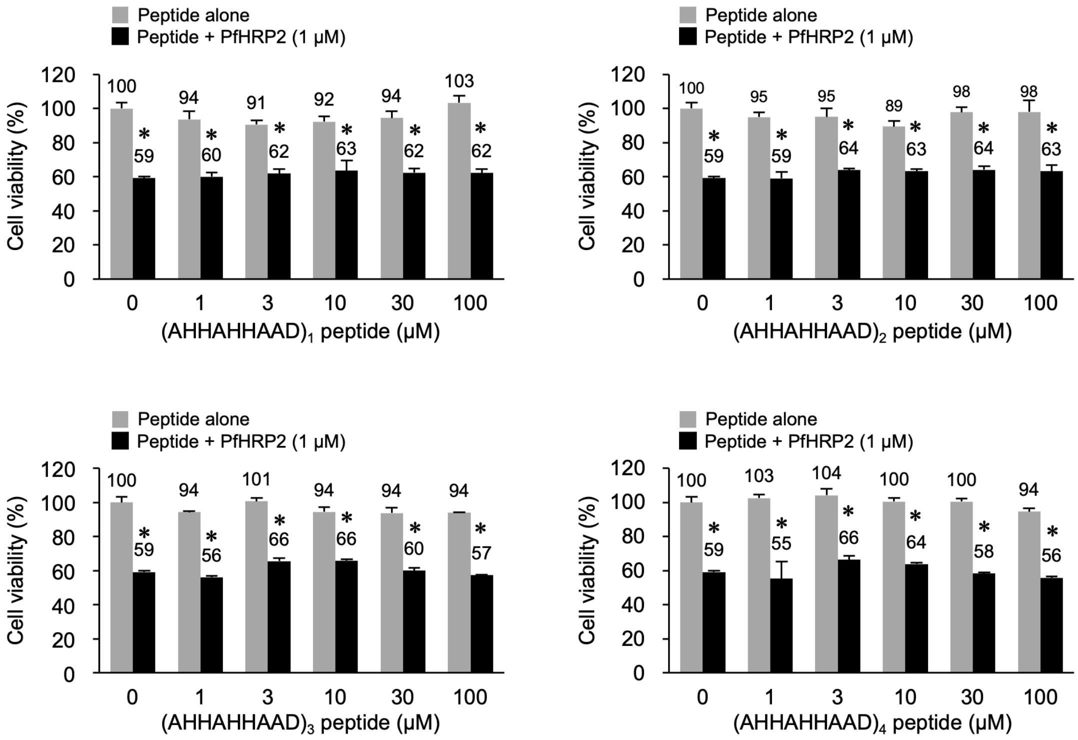


**Fig. S3.** Negligible effects of repetitive AHHAHHAAD peptides on PfHRP2 cytotoxicity in HT1080 cells. PfHRP2 cytotoxicity was determined in the presence of partial peptides of PfHRP2. HT1080 cells were treated with PfHRP2 (1 μM) at 37°C for 24 h in the presence of partial peptides (AHHAHHAAD$_1$, AHHAHHAAD$_2$, AHHAHHAAD$_3$, and AHHAHHAAD$_4$ peptides). Relative cell viability was determined using the Cell Counting Kit-8 reaction solution. Means and SD are shown (*$p$ < 0.05). n.s. indicates a nonsignificant difference.

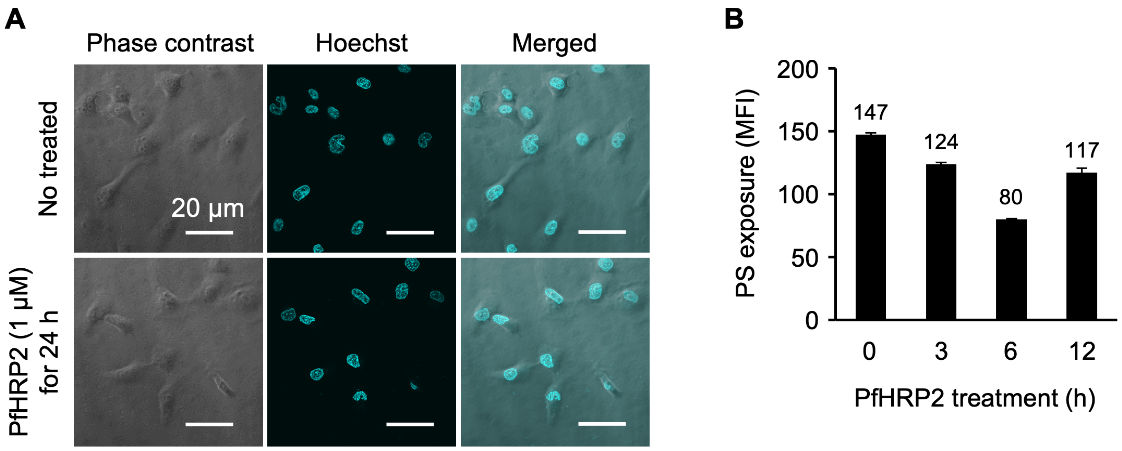


**Fig. S4.** Apoptosis-independent PfHRP2 cytotoxicity in HT1080 cells. (*A*) Morphology of HT1080 cells treated with PfHRP2. The HT1080 cells were incubated under the condition that PfHRP2 shows efficient cytotoxicity (1 µM of PfHRP2 at 37°C for 24 h). Cellular morphology was observed using a confocal laser scanning microscope. Cyan fluorescence indicates nuclei stained with Hoechst 33258. Scale bars: 20 µm. (*B*) Phosphatidylserine (PS) exposure on the cellular membrane of HT1080 cells treated with PfHRP2. HT1080 cells were treated with PfHRP2 (1 µM) at 37°C for 3–12 h. PS exposure, known as an early apoptosis marker, was detected using an Annexin V assay kit (BioVision, CA, USA) and evaluated according to mean fluorescent intensity (MFI) via flow cytometric analysis. Means and SD are shown.

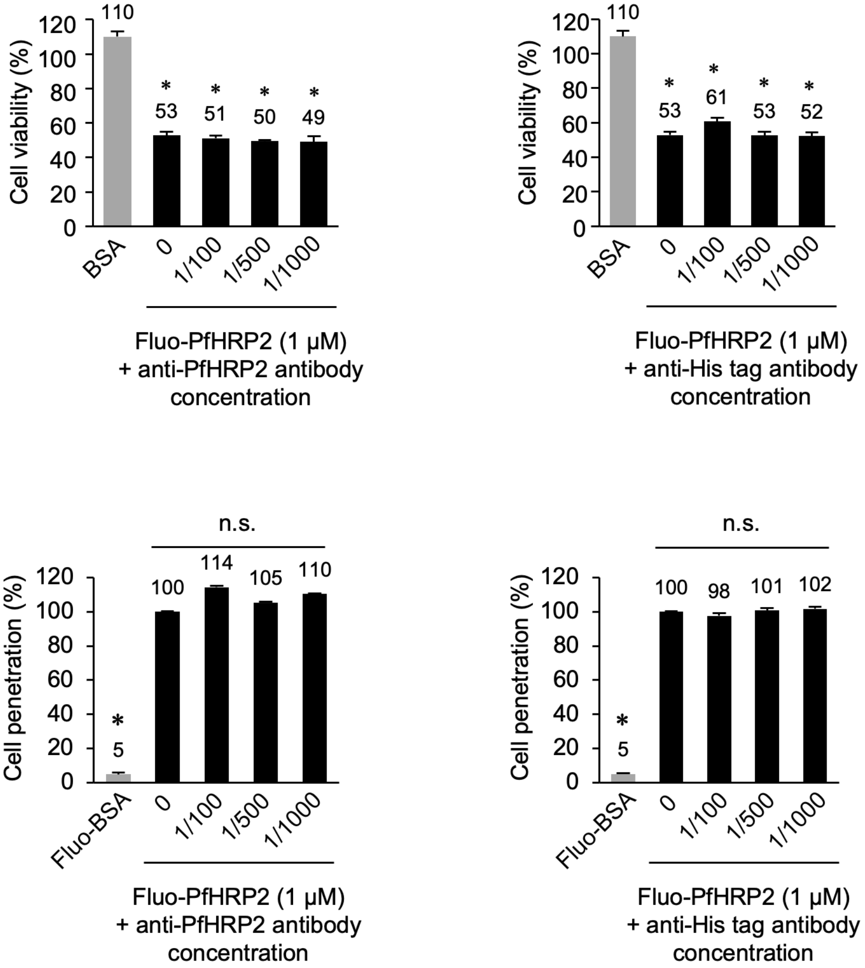


**Fig. S5.** Negligible effects of antibodies on PfHRP2 in HT1080 cells. Cytotoxicity and cell penetration of PfHRP2 were determined in the presence of antibodies. HT1080 cells were treated with PfHRP2 (1 μM) at 37°C for 24 h in the presence of an anti-PfHRP2 antibody (x1/100–x1/1000) or anti-His tag antibody (x1/100–x1/1000). Relative cell viability was determined using the Cell Counting Kit-8 reaction solution. Conversely, for cell penetration, HT1080 cells were treated with Fluo-PfHRP2 (1 μM) at 37°C for 3 h in the presence of anti-PfHRP2 antibody (x1/100–x1/1000) or anti-His tag antibody (x1/100–x1/1000). Relative cell penetration of proteins was evaluated by MFI in the flow cytometric analysis. BSA and Fluo-BSA were used as negative controls. Means and SD are shown (*$p < 0.05$). n.s. indicates a nonsignificant difference.

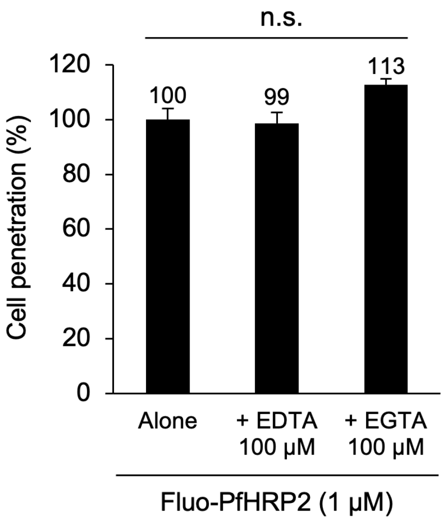


**Fig. S6.** Cellular uptake of PfHRP2 in the presence of EDTA/EGTA. HT1080 cells were treated with Fluo-PfHRP2 (1 μM) at 37°C for 3 h in the presence of EDTA (100 μM) or EGTA (100 μM). Relative cell penetration of PfHRP2 was evaluated by mean fluorescent intensity (MFI) in flow cytometric analysis. Means and SD are shown. n.s. indicates a nonsignificant difference.

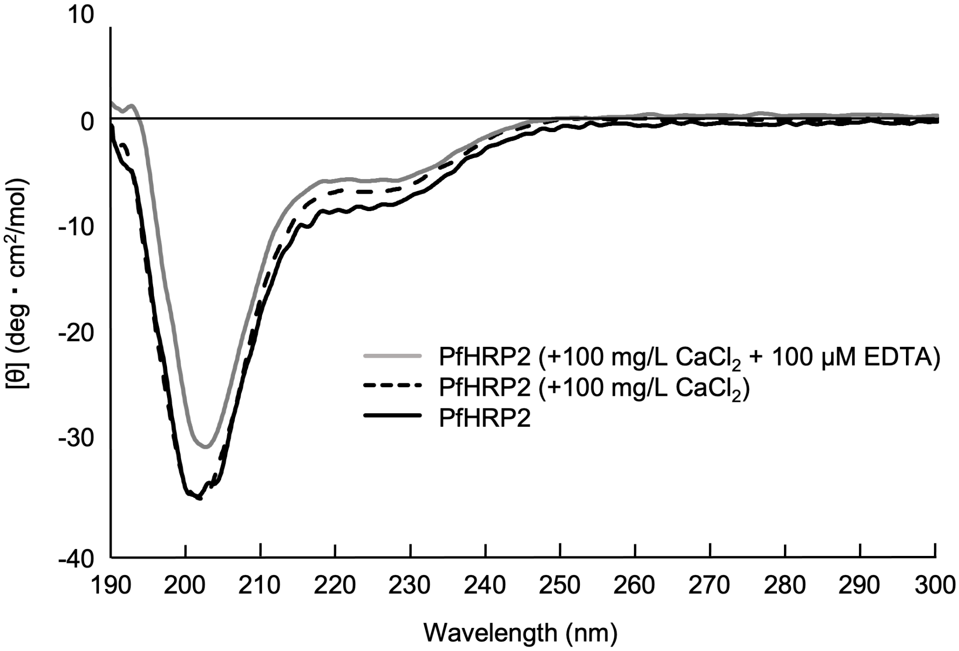


**Fig. S7.** Circular dichroism (CD) spectrums of PfHRP2 in the presence of $CaCl_2$ and EDTA. PfHRP2 (1 μM) was incubated with or without $CaCl_2$ (100 mg/L) and EDTA (100 μM) in 20-mM phosphate buffer (pH 7.4) at 37°C for 1 h. Additionally, the CD spectrums of PfHRP2 with/without $CaCl_2$ and EDTA were measured using a 1-mm path length cell at 37°C on a Jasco J-820 CD spectrophotometer.

**Table S1.** Components of $CaCl_2$-modified medium.

| **Medium components (mg/L)** | | | | |
|---|---|---|---|---|
| **$CaCl_2$** | **0** | **25** | **50** | **100** |
| $MgSO_4$ | 100 | | | |
| KCl | 400 | | | |
| NaCl | 6,800 | | | |
| $NaHCO_3$ | 2,200 | | | |
| $NaH_2PO_4$ (anhyd.) | 121.74 | | | |
| L-Arginine HCl | 126 | | | |
| L-Cystine | 23.78 | | | |
| L-Glutamine | 292 | | | |
| L-Histidine HCl $H_2O$ | 42 | | | |
| L-Isoleucine | 52 | | | |
| L-Leucine | 52 | | | |
| L-Lysine HCl | 72.5 | | | |
| L-Methionine | 15 | | | |
| L-Phenylalanine | 32 | | | |
| L-Threonine | 48 | | | |
| L-Tryptophan | 10 | | | |
| L-Tyrosine | 36 | | | |
| L-Valine | 46 | | | |
| D-Pantothenic acid | 1 | | | |
| Choline Chloride | 1 | | | |
| Folic Acid | 1 | | | |
| i-Inositol | 2 | | | |
| Niacinamide | 1 | | | |
| Pyridoxal HCl | 1 | | | |
| Riboflavin | 0.1 | | | |
| Thiamine HCl | 1 | | | |
| D-Glucose | 1,000 | | | |
